# Optimizing functional groups in ecosystem models: Case study of the Great Barrier Reef

**DOI:** 10.1101/655076

**Authors:** Vanessa Haller-Bull, Elena Rovenskaya

**Affiliations:** School of Mathematical Sciences, Queensland University of Technology, 2 George St, Brisbane, 4000, Qld, Australia; International Institute of Applied System Analysis (IIASA), Schlossplatz 1, Laxenburg, 2361, Austria; Faculty of Computational Mathematics and Cybernetics, Lomonosov Moscow State University, Leninskie Gory, 1(52), GSP-1, Moscow 119991, Russia

**Keywords:** structural uncertainty, resolution, node aggregation, ecosystem models, foodweb model, coral reefs

## Abstract

Uncertainty is inherent in ecosystem modelling, however its effects on modelling results are often poorly understood or ignored. This study addresses the issue of structural uncertainty or, more specifically, model resolution and its impact on the analysis of ecosystem vulnerability to threats. While guidelines for node assignments exist, they are not underlined with quantitative analysis. Different resolutions of a coral reef network are investigated by comparing the simulated network dynamics over time in various threat scenarios. We demonstrate that the error between a higher-resolution and a lower-resolution models increases, first slowly then rapidly with increased degree of node aggregation. This informs the choice of an optimal model resolution whereby a finer level of a food web representation yields only minimal additional accuracy, while increasing computational cost substantially. Furthermore, our analysis shows that species biomass ratio and the production ratio are important parameters to guide node aggregation to minimize the error.

## 1 Introduction

Simplification of reality and related uncertainty is unavoidable in any applied research aiming to support decision making; what influences the quality of analysis is how that uncertainty is incorporated into management decision processes. This issue was highlighted by Ludwig et al. (1993), who said: “effective policies are possible under conditions of uncertainty, but they must take uncertainty into account”. Walker et al. (2003) defines uncertainty as “any deviation from the unachievable idea of completely deterministic knowledge of the relevant system”. Another way of thinking about uncertainty is that “uncertainty reflects the probability that a particular estimate, piece of advice, or management action may be incorrect” (Lek, 2007). While these definitions describe uncertainty well, they do not differentiate between different types of uncertainty.

Uncertainty can arise at different instances in a modelling process. For the purpose of this paper, we will concentrate on two types of uncertainty. *Parameter uncertainty* is defined as the difference between the true value of a parameter and the mean value estimated using the data available and statistical techniques (Skinner et al., 2014). *Structural uncertainty* refers to a mismatch between the simplified mathematical equations of a model and the true complex ecological relationship observed in situ (Refsgaard et al., 2006). One type of structural uncertainty, which is the one we focus on in this study, is model resolution. In an ecological network, resolution refers to the number of nodes in the network. A node within the network represents an ecological unit that can be at different aggregation levels incorporating one or more species. Generally, it is believed that low-complexity models reduce parameter uncertainty by reducing the number of parameters, while more complex models reduce structural uncertainty since they more closely describe the natural system (Iwasa et al., 1987). This leads to a hump shape between the level of complexity and the accuracy, with medium complexity models often performing best (Håkanson, 1995, Costanza and Sklar, 1985, Jester, 1977).

Even though it has been acknowledged for a long time that different types of uncertainties are crucial, their role is often not thoroughly understood, especially in complex models (Milner-Gulland and Shea, 2017, Link et al., 2012). Weijerman et al. (2015) completed an extensive review of ecosystem models and of the 27 ecosystem models reviewed only one addressed structural uncertainty as well as parameter uncertainty.

Ecosystems are complex due to the combination of multifaceted species interactions, which are often nonlinear, which may result in multiple equilibria (Gordon, 2007, McClanahan et al., 2009). Furthermore, ecosystem models are often faced with many threat adding to their complexity. The management of these threats on the systems is one of the main modelling goals. Ecosystem models always involve high uncertainties, especially when attempting to predict the effects of interventions and management actions (Costanza et al., 1993, Hill et al., 2007). If uncertainty is larger than believed, the results are more likely to be misleading, and are therefore more likely to generate an inefficient or incorrect management decision (Weijerman et al., 2015). Lek’s definition of uncertainty emphasizes this possibility of decision-makers being misled by models.

Ecosystem models are usually created at a functional group level to reduce the number of nodes and therefore parameters (Fulton et al., 2003). A functional group refers to a group of species that are assumed to be so similar in a defined set of characteristics that they can be investigated as one unit. In the literature, species are commonly grouped according to their trophic status and diet (e.g., herbivore, or detritivore) (Stoddart, 1969). In ecosystem modelling a functional group is often used as the basis of one node: Instead of treating each species as different and assigning a different dynamic equation and parameters to each species, the functional group is assumed to be homogenous enough to be represented by a single equation. Reducing resolution by considering functional groups only of course reduces the total number of nodes in the system, which consequently, reduces the complexity and the number of parameters that needs to be estimated. Guidelines about which species to group together have existed for a long time (Gardner and Ashby, 1970, Wiegert, 1975, O’Neill, 1975, Cale Jr and Odell, 1980), however, there has been no formal investigation of how much uncertainty is caused by the introduction of functional groups suggested in the literature or grouped according to these guidelines. Thus, we lack an understanding of the magnitude and even distribution of structural uncertainty in the threat response of ecosystems (Bellwood and Fulton, 2008).

Ecopath with Ecosim (Colléter et al., 2015) is a common modelling framework used to analyse marine systems, especially in the context of fisheries management. Ecopath is based on the estimation of biomasses and food consumption to create a mass-balanced food web. It has been reviewed and extended on over the past 40 years to enable dynamic simulations (Ecosim) and spatial analysis (Ecospace) (Christensen and Walters, 2004). Since its inception, over 800 studies have used it to investigate questions related to fisheries management.

In this study, we develop and apply a new approach of varying the resolution of an ecological network while simulating the food web response to a species degradation. The aim of this study is threefold:

1. We want to show that uncertainty introduced due to lowering the system resolution even slightly can be substantial and should not be neglected.
2. We want to propose a basis for choosing an optimal resolution to balance parameter and structural uncertainty.
3. We want to suggest ways to improve the techniques used to group species in ecosystem models while also generalising these techniques for other types of networks.

## 2 Methods

The goal of this study is to understand how the ecosystem resolution affects the model’s predictive power in what concerns the effects of species degradation onto the entire ecosystem. To do so, we will examine a number of “threat” scenarios, in which a fraction of a biomass of a focal species is removed giving rise to changes in biomasses of other species because of feeding relationships. We will evaluate the variation between threat impacts modelled based on a network at species level versus a network at a functional group level, which has a coarser resolution. To calculate errors, it is assumed that the high-resolution model reflects nature closely and thus the error between the high-and low-resolution models can be used as an estimate of the actual uncertainty. As the resolution of a model is a continuous characteristic, in this study, nodes within a chosen functional group are being merged, one by one, to “imitate” a step-by-step formation of the functional group. This enables a comparison of the response dynamics under the same input (a particular threat scenario) and an estimation of the difference that each particular merger entails. On this basis, a thorough understanding of the changes in the error and, consequently, the uncertainty with a reduction in resolution can be obtained.

Naturally, one expects that a model should become more “precise” with a finer resolution. However, it comes at a cost of computing power that is required. In this study we refer to an “optimum” in terms of good precision without an explosion of computational power.

### 2.1 Ecosystem simulation model and calibration

The dynamic ecosystem model used in this study is based on a network of interacting nodes that represent a single or a group of species. The interactions considered between the nodes resemble predation, i.e., the consumption of biomass of one node by another. This means that the connection between two nodes indicates organic matter being transported from one node to the other and travelling generally up the food chain. Organic matter can be lost from the living nodes to the external environment through energy spent in respiration. Egestion and or other mortality that add to the detritus close the loop from high to low trophic levels of the food chain.

The network model used for this study is based on the Ecopath with Ecosim toolbox. We use Ecopath to calibrate our ecosystem network model and then we adopt Ecosim equations to simulate the ecosystem dynamics in response to a shock. We re-code the model in MatLab for convenience of running simulations. In what follows, we present a general approach towards achieving the goal of this study as discussed above, and then we apply it to Great Barrier Reef as a case study.

First, we describe the modelling framework. Ecopath is a static mass-balanced ecosystem model describing the network through linear equations that connect biomasses, diet preferences, mortalities and productions in an ecosystem. The static equations of Ecopath can then be extended to differential (difference) equations to allow for dynamic manipulations of the system (Ecosim extension).

Ecopath is based on two master equations. The first master equation (1) focuses on the mass balance within each nodes, while the second master equation (2) describes the mass balance between nodes; the first master equation is as follows

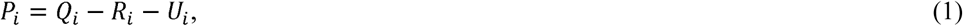

where *P*_*i*_ represents the total biomass production of node *i*(*t km*^−2^*year*^−1^) and *Q*_*i*_ represents the total biomass consumption of node *i*(*t km*^−2^*year*^−1^), *R*_*i*_ represents the respiration of node *i* (*t km*^−2^*year*^−1^), which is a loss of a part of biomass to the environment and *U*_*i*_ represents the amount of unassimilated food for each node, *i* (*t km*^−2^*year*^−1^) for all *i*=1,…*n* with, being the number of nodes in the ecosystem network. Unassimilated food refers to the amount of biomass lost through excretion, i.e., the amount of biomass input into the lowest trophic level node called detritus.

The second master equation unfolds the node production of all nodes (besides detritus) that consists of the amount of its biomass consumed by other nodes plus the amount of biomass that emigrates and the amount of biomass that is lost due to natural mortality, here called other mortality or excretion. The flow from one node to another is assumed to be proportional to the total consumption of the node-recipient, which is sometimes called top-down or recipient control:

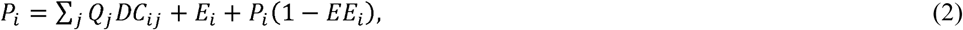

where *DC*_*i,j*_= proportion of the consumption by predator *j* that is made up of prey *i* (unitless), *E*_*i*_= net emigration (emigration – immigration; *t km*^−2^*year*^−1^); and *EE*_*i*_ = ectotrophic efficiency (unitless). The ecotrophic efficiency is the proportion of production that is passed onto the next trophic level. This parameter is smaller than one due to natural mortality.

This formulation of the second master equation assumes that there is no biomass accumulation (in Ecopath notations *BA*_*i*_=0) to ensure a steady state and as a baseline it also assumes that there was no fishing (*Y*_*i*_=0).

Combining (1) and (2) one obtains

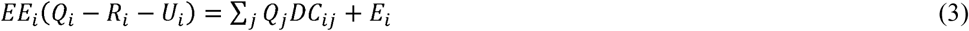

which means that the surviving production of one group (the left-hand side of this equation) is equal to the consumption of that group by all predators and the biomass that is leaving through emigration (see Figure 1).

**Figure 1.**
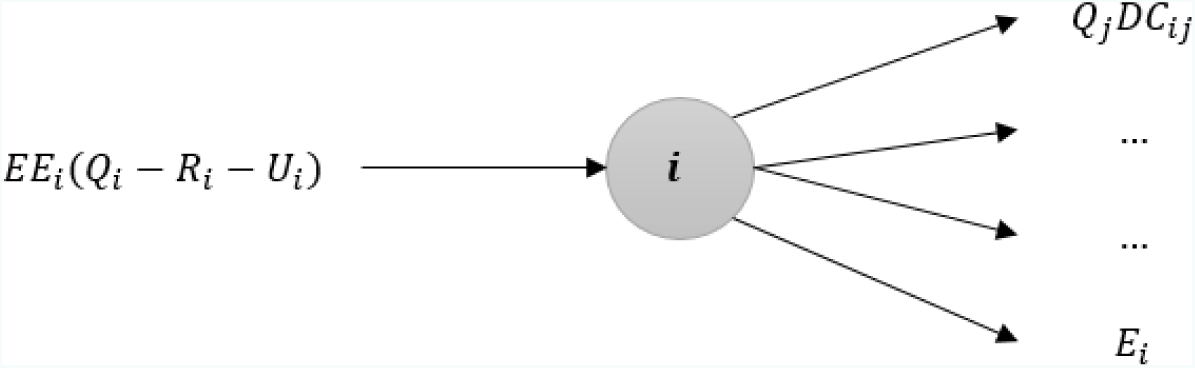
Visualisation of equation (3), the mass-balance of a single node within the network

The data requirements for Ecopath encompass at a minimum the input of parameters *DC*_*ij*_ and *E*_*i*_, as well as any three out of four variables *P*_*i*_,*Q*_*i*_,*B*_*i*_ and *EE*_*i*_ for each node in the model. The missing parameters are then estimated using the mass-balance equations (1) and (2); with *T*_*i*_=0 for all *i*=1,….*n*, this process is also called balancing the Ecopath model.

For some nodes we might have information on all four parameters. However, since they have often been estimated during different studies and any such parameters contain uncertainty, we often cannot reach a mass-balance directly by just calculating the remaining unknown parameters (which are fewer than *n*). In this case, we conduct another step called the model calibration. For the model calibration, i.e. to find “free” parameters that ensure mass-balance, all parameters are varied according to the parameter uncertainty underlying their estimates until a mass-balance is reached. For example, the production of a species might have been investigated and confirmed by several studies, so the parameter uncertainty is low and the bound in which it is varied for calibration might be set at 10%. On the other hand, a production estimate of a different species might not be available for the study region and needs to be inferred from a different region. This would result in a higher parameter uncertainty and hence a variation bound of 20% would be used. Rules to identify these uncertainty bounds have been established and are called pedigrees within the Ecopath software (Christensen et al., 2005). In practice this means that we can often find several possible parameterizations that ensure mass-balance. To identify robust conclusions for a study utilising these models several/ all of these parameterizations should be evaluated. We elaborate more on how we used them in this study in section 2.5.

Equation (2) motivates a differential equation in which species biomass evolves over time as a response to external shocks and pressures, such as harvesting. Let *B*_*i*_ be the biomass of a group *i*. From here on out we no longer consider the total production (*P*_*i*_) or consumption (*Q*_*i*_), but rather the production (*p*_*i*_) and consumption (*q*_*i*_) per unit of biomass. Since (2) has the assumption of a steady state, we can rearrange it to refer to the population change of zero as follows

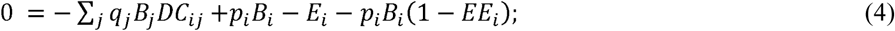

or generally as change in population size over time

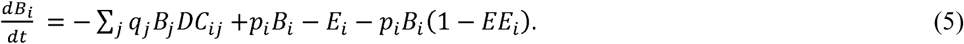

Furthermore, we consider that immigration *I*_*i*_ into the group is independent of the population sizes within the system, while emigration *e*_*i*_ out from the group depends on the population size of group *i,,* due to the effect of crowding, which means

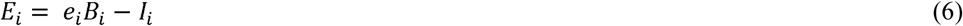

We denote

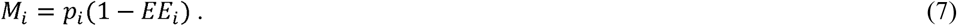

which refers to the non-predation mortality. So, all together we obtain

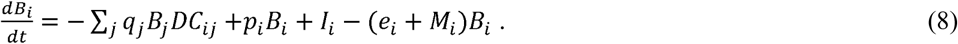

In the literature, equation (8) is often described in a more general form

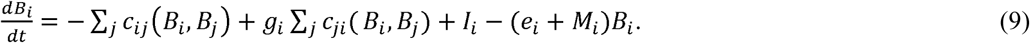

Equations (9) has the advantage that *c*_*ij*_(*B*_*i*_,*B*_*j*_) can be chosen to represent bottom-up, top-down or mixed control. In this study, we investigate bottom-up control due to its inherent stability (Hearon, 1963) and consequently, assign

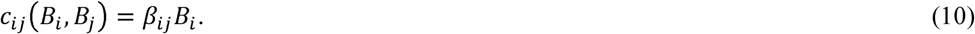

Since each flow from *i* to *j* needs to be consistent between the static and the dynamic approach we define

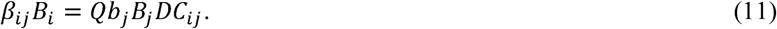

Consequently, **β** _*ij*_ is defined as

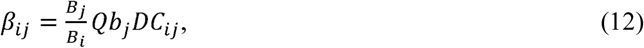

where all entries in formula (12) are to be taken from the model calibration results.

Thus, we have

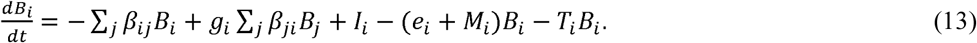

where *g*_*i*_ is the growth efficiency, which is a ratio between the production and consumption (unitless) and *β* _*ij*_ represent elements of the interaction matrix (Christensen et al., 2005)(Christensen, Walters and Pauly 2005)(Christensen, et al., 2005)(Christensen et al., 2005)(Christensen et al., 2005)(Christensen et al., 2005)(Christensen et al., 2005)(Christensen et al., 2005)(Christensen et al., 2005). In (13), we also include threats (last term in the right-hand side) with *T*_*i*_ being the proportion of the total biomass removed by threat *i* in Ecopath usually only considered as fishing (*t km*^−2^*year*^−1^). Most threats remove part of a population so 0<*T*_*i*_ <1, however it could also be −1<*T*_*i*_ <0 in case of a threat that increases an unwanted population, for example the increase of algae,

In the dynamic simulations, we initiate system (13) using initial values *B*_*i,t*=0_ equal to the steady-state values *B*_*i*_ calculated using Ecopath. Emigration and immigration are also taken from Ecopath. As the dynamics is affected by a threat, the system is moving away from the initial steady state corresponding to an equilibrium in the absence of external disturbance. Due to the donor-control assumption, the dynamics converges to a new steady state.

### 2.2 Case study system and data

The Great Barrier Reef (GBR) is the biggest reef system in the world. It is currently under a severe threat due to global warming and the resulting bleaching. One reef within this system is Rib reef (lat:-18.48, long:146.88). It is located on the mid-shelf in the central Great Barrier. Due to its location, Rib reef is a good template for other reefs within the system. Rib reef was previously investigated in a fisheries management assessment study using Ecopath (Tudman, 2001). This study used an Ecosim network with 25 nodes representing functional groups. However, data in this paper is simply a collection of studies and as such it is available from the original published sources at the species level for most fish species, allowing for a more complex network of *n*=206 nodes. This makes Rib reef an ideal case study for evaluating the consequences of merging species into functional groups for modelling and management. Note that in this case only fish species are identified at a species level, while other functional groups (total of 15) are kept at a functional group level to reduce the data requirements and the model complexity and consequently to keep computational time feasible (in the absence of these limitations the model resolution could be increase to a few thousand nodes). When interpreting our results here, this limitation needs to be remembered, i.e., the actual error due to coarse model resolution is likely to be larger than estimated here.

Following Tudman (2001), we adopt input parameters *P*_*i*_,*Q*_*i*_,*B*_*i*_ for all *i*=1,…*n* and the net emigration *E*_*i*_ as zero. Parameters *DC*_*ij*_ for all *i,j*=1,…*n* are downloaded directly from Fishbase (Froese and Pauly, 2017), as these parameters were only given at a functional group level in Tudman (2001). Assuming that the ecosystem is initially in the steady state, using the available data for *P*_*i*_,*Q*_*i*_,*B*_*i*_ and *DC*_*ij*_ we derive the missing parameters *EE*_*i*_. This is possible if at least two parameters of each node are known. In this way, we fully calibrate our dynamic model, the resulting parameters for the following simulations can be found in the supplementary materials (S1).

### 2.3 Threat implementation and threat scenarios

We model threats as a fixed proportional reduction of biomass of a node at each time step. This is a common way to represent fishing pressures in ecosystem models, and it can also represent other threats such as bleaching or nitrification via reducing or increasing the biomass of a lower trophic level.

We implement a total of six threat scenarios in this study to prototype a range of alternatives from fishing on a variety of target species to coral bleaching and increased nutrition choosing those scenarios which represent most typical and critical threats for coral reefs. Importantly, these scenarios represent threats impacting a variety of trophic levels from top predators to primary producers and consequently, enable a detection of potential differences in terms of uncertainty spread unevenly across trophic levels. However, it is important to keep in mind that the selected scenarios are theoretical and not directly based on data. The full list of scenarios is as follows:

1. A reduction of the biomass of pelagic fish (fishing on a high trophic level)
2. A reduction of the biomass of coral trout (fishing on a high trophic level)
3. A reduction of the biomass of herbivores (fishing on a medium trophic level)
4. A reduction of the biomass of sharks (fishing on a high trophic level)
5. A reduction of the biomass of coral (indicative of bleaching)
6. An increase of the biomass of algae (indicative of increased nutrients).

All of these scenarios are introduced at different intensities ranging from 10 to 90% removal/introduction rate, i.e., *T*_impacted node_=0.1,0.2,…,0,9 (per year).

### 2.4 Merging nodes and error calculation

Merging nodes refers to pooling of biomasses of two nodes (here species or functional groups) into one node and corresponding uniting/reassignment of incoming and outgoing flows. The following describes the routine of consecutive mergers we use to evaluate the error for each of the 54 scenarios (six different threats and nine intensities per threat). In the first instance, we focus on merging nodes within each functional group from the original model (Tudman, 2001) individually, meaning that all nodes outside the functional group in focus remain at species level. Then we merge the functional groups consecutively to fully recreate the original 25-node model.

For each of the eight functional groups in the original model we run an “experiment”. Each experiment consists of a number of steps equal to the initial size of the functional group in terms of the number of nodes. At each experiment step, we reduce the number of nodes from the functional group by one from *j* to *j* − 1. To do so, we implement all possible mergers of two nodes within this functional group into one, total 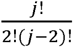 mergers. For every merger at a given experiment step, we consecutively run all six threat scenarios with nine intensity levels, total 54 scenarios. For each scenario, we simulate the ecosystem dynamics using (13) and initial conditions from Ecopath for up to 500 time steps (which is equivalent to 5 years) and compare it with the dynamics corresponding to the case without any mergers. To quantify the comparison we use the total relative error as follows

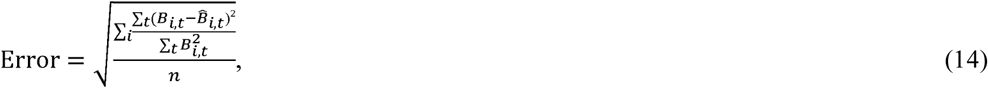

where 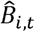 refers to the merged biomass of a node and *B*_*i, t*_ refers to the unmerged biomass of the two merged nodes added together after the simulations; *n* here is the size of the considered ecosystem model version.

Since five years is a commonly used timeframe for management decision making, it is chosen as the length for the simulations.

Actually, due to the linearity of the model equations, for a given threat (including none) the initial (or any given) steady state of the full system can be reproduced exactly by the reduced system (see supplementary materials S2) by a choice of coefficients *DC*_*ij*_, as well as *B*_*i*_,*P*_*i*_ and *Q*_*i*_ in the reduced system. But modellers and decision makers are interested in models which are able to process a variety of possible threats, hence we use a metric (14) to evaluate the “goodness” of a merger averaging over all nodes in the system. Also (14) focuses on the transient path instead of just the eventual steady state.

At each experiment step, we save the “best” merger and a corresponding network, resulting in the lowest error (14). We use this “optimal” network configuration as a starting point for the next experiment step. Experiments continue until the considered functional group becomes a single node.

### 2.5 Parameter and structural uncertainty

In a study that would like to estimate the response of this coral reef to a given threat, the stepwise aggregation that we have done so far is of not much use. Managers would be interested in the total amount of uncertainty around the population size estimates that we are provided with a given model. This situation is what we are exploring in this section. The total amount of uncertainty in the estimates is partly due to parameter uncertainty and party due to the structural uncertainty. Here, we are comparing how much uncertainty we would assign depending on if we consider both the structural and the parameter uncertainty or only one of them as it is often done in previous studies (Weijerman et al., 2013).

Parameter uncertainty can be easily estimated by simulating the system response with different parameter sets. As described in the previous section 2.1 we calibrate the model with different variations of the original parameters to create 10 separate parameter sets for the system. The parameter sets are then each simulated over 500 time steps (5 years) for each scenario and intensity. The resulting responses are compared to the response of original parameter set using the same error estimates as for the structural uncertainty described in section 2.4. For these error estimates (10 *parameter sets* * 6 *scenarios* *9 *intensities*=540 *estimates*) (we then calculate the mean and standard error.

Structural uncertainty here is based on errors that are calculated similar to the previous methods in section 2.4. The only difference is that instead of calculating the error at each merger, we create three resolutions and only compare the error between these. The three resolutions are as followed (S3): First, we have the full resolution model (assumed to have no structural error) with 206 nodes. Second, we have the medium resolution model identified as the optimum from the previous sections with 49 nodes. Third, we have the low resolution model based on the functional group in the original study by Tudman. This results in 54 error estimates (6 *scenarios* *9 *intensities*) for each resolution which are then averaged to calculate the structural uncertainty.

Now we can combine both of these uncertainties by creating 10 parameter sets for each resolution and comparing the errors across all of them (10 *parameter sets* * 6 *scenarios* *9 *intensities*=540 *estimates for each resolution.*

### 2.6 Regression tree analysis

In order to evaluate the error introduced through a merger of two nodes according to the characteristics of these nodes we use regression analysis. Characteristics of interest are related to parameters of each node, i.e., biomass, production, or specific to each combination of nodes in conjunction with the threat scenario, i.e., the difference in trophic level between nodes merged and nodes experiencing a threat (Table 1). We use the bagged regression tree analysis since input data, predictors and responses typically have different non-normal distributions. A regression tree analysis is a supervised (ie. there is a response variable) machine learning algorithm. At each step, the algorithm splits the samples according to one of the predictors to form a more homogenous group of the response. This means that if we know the value for each predictor we can follow all of the splits and predict the likely response. An extension of this is used in this study by growing several trees (200) for more reliable results. In the bagged regression tree analysis we grow each regression tree while utilising all predictors but subsampling data from the initial sample (Prasad et al., 2006). Those trees are then combined in an ensemble to give more reliable predictions. We performed an analysis with replacement i.e. each subsample is placed back into the full sample after growing the tree and a new subsample is taken for the next tree. This provides an opportunity for internal validation at each tree level using the mean squared error (MSE) of predictions. Compared to other methods of growing ensembles of trees, bagged trees largely prevent overfitting. Another great feature about the bagged tree analysis is that it can calculate surrogate splits. At each split, when the tree is grown, the algorithm determines the next best split according to the MSE, this split is then known to be the surrogate split. Overall, this procedure enables comparing the optimal split (the split chosen by the algorithm to grow the tree due to the lowest MSE) and the surrogate splits for all variables. The variable association is then measured according to how different the MSE between the optimal and the surrogate split is. Variables that can be easily replaced by their surrogate splits are less important and could be excluded. Additionally, we also calculate the overall variable importance, which is the total reduction of the MSE that is due to splits based on that variable (Prasad et al., 2006). This means that if the variable importance is high, we the splits based on that particular predictor are have a large influence on the reduction of the MSE achieved by the analysis. Variables with a low importance only reduce the MSE a little, so could potentially be excluded from the analysis without much loss in overall performance. The variable importance calculated here is relative within the analysis, i.e. the magnitude of the importance can only be compared between variables of the same analysis not across different models.

**Table 1:**
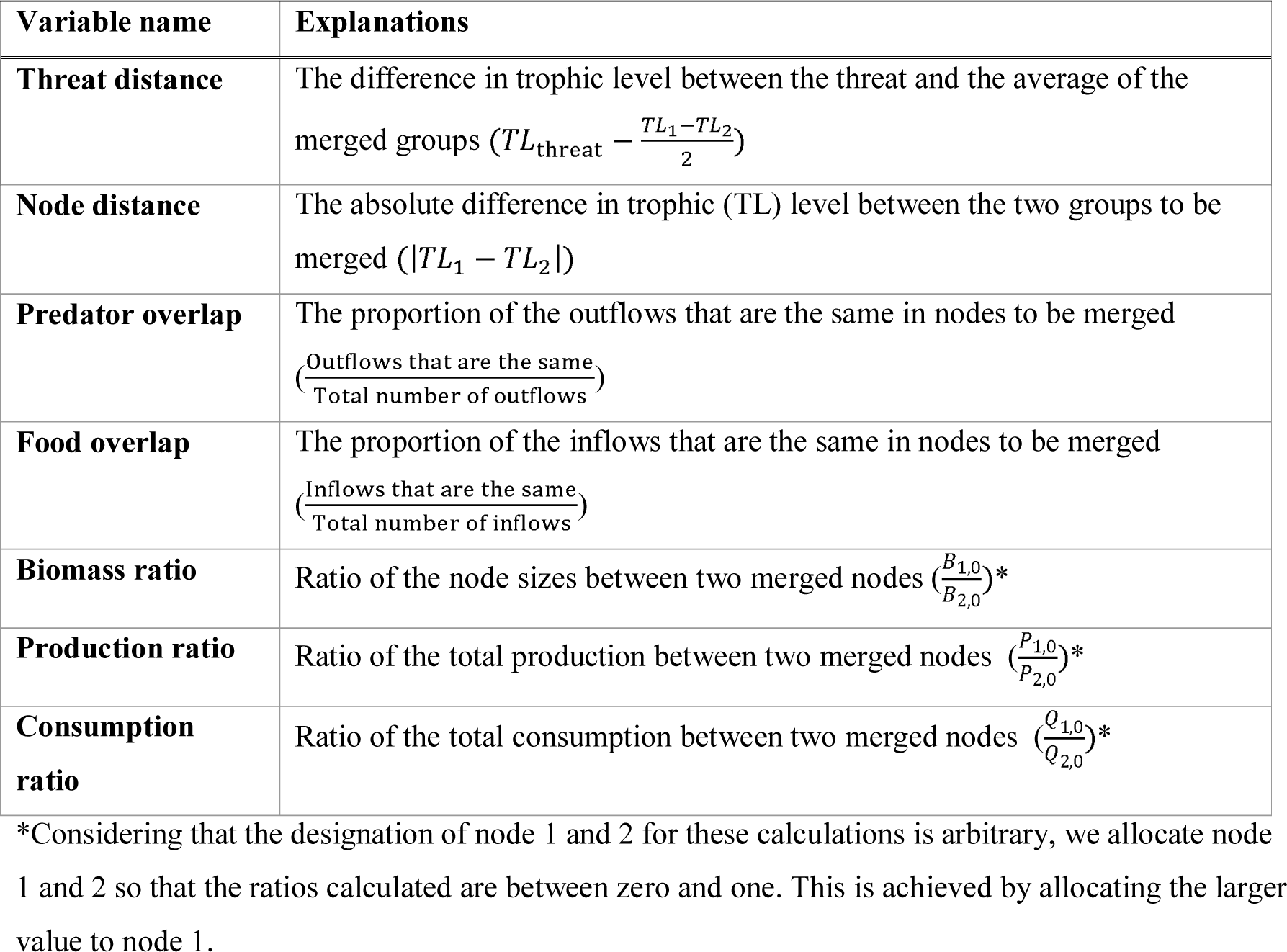
Predictors utilised in the bagged tress analysis

## Results

### 3.1 Error due to model coarse-graining

The error estimates after merging show a clear pattern, the error estimates start low then first slowly then rapidly increase. However we are also interested in differentiating between the maximum and minimum error at each merger. Since this represents the variation of errors that can occur depending on if we merge two similar or two very different species or nodes. For most scenarios, the minimum error is rather small when only few nodes are merged, however after a “tipping point” when a certain “critical mass” of nodes is merged, it increases rapidly. The maximum error on the other hand starts relatively high and remains mostly stable with just a small increase at the final mergers. This also means that we can reduce the error when only merging a few species, however only if we are merging the correct species. Finally, the spread of errors across the 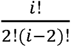 possible networks at each merger is not even, but rather clustered. This clustering can aid the identification of the nodes (here species) that should or should not be merged to avoid large losses of accuracy.

Figures 2 and 3 illustrate the typical patterns we observed in these experiments. Figure 2 depicts errors in one (of six) threat scenario with fixed intensity (0.5) for all eight functional groups. Figure 3 illustrates the effects of all six threat scenarios with fixed intensity (0.5) on one selected functional group (lutjanids). Both figures represents the tendencies concerning the minimum and maximum errors described above from two different perspectives. The graphs of all functional groups across all six scenarios and intensities can be found in the supplementary materials (S4).

**Figure 2.**
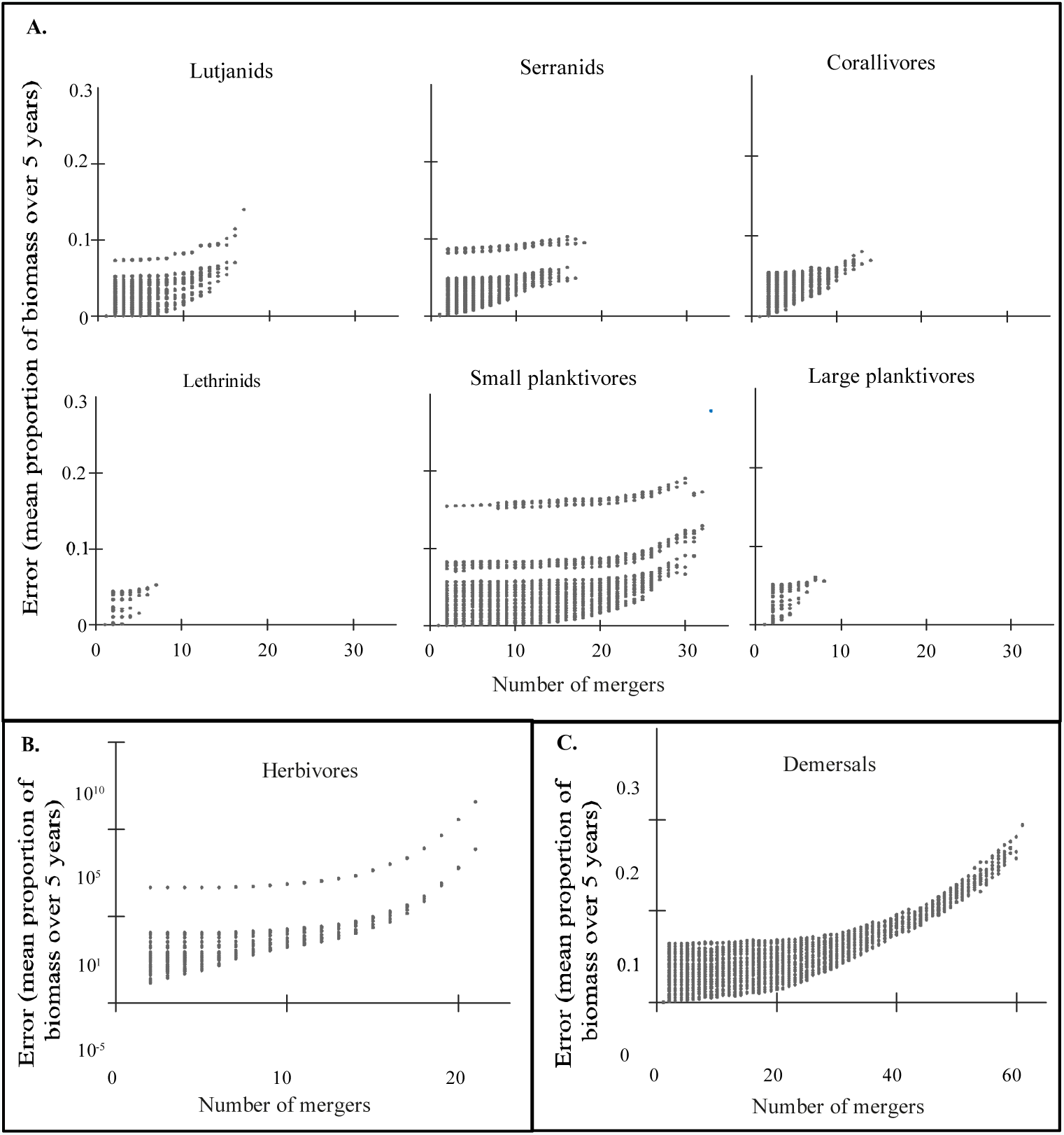
Example of results: Error estimates for all eight functional groups under threat scenario 1) in which the biomass of pelagic fish is reduced with intensity 0.5. Panel A compares functional groups lutjanids, serranids, corallivores, lethrinids, small and large planktivores. Panel B shows herbivores and panel C the functional group other demersals. Each point represents one possible merger of two nodes originating from the model identified with the lowest error in the previous mergers.

**Figure 3.**
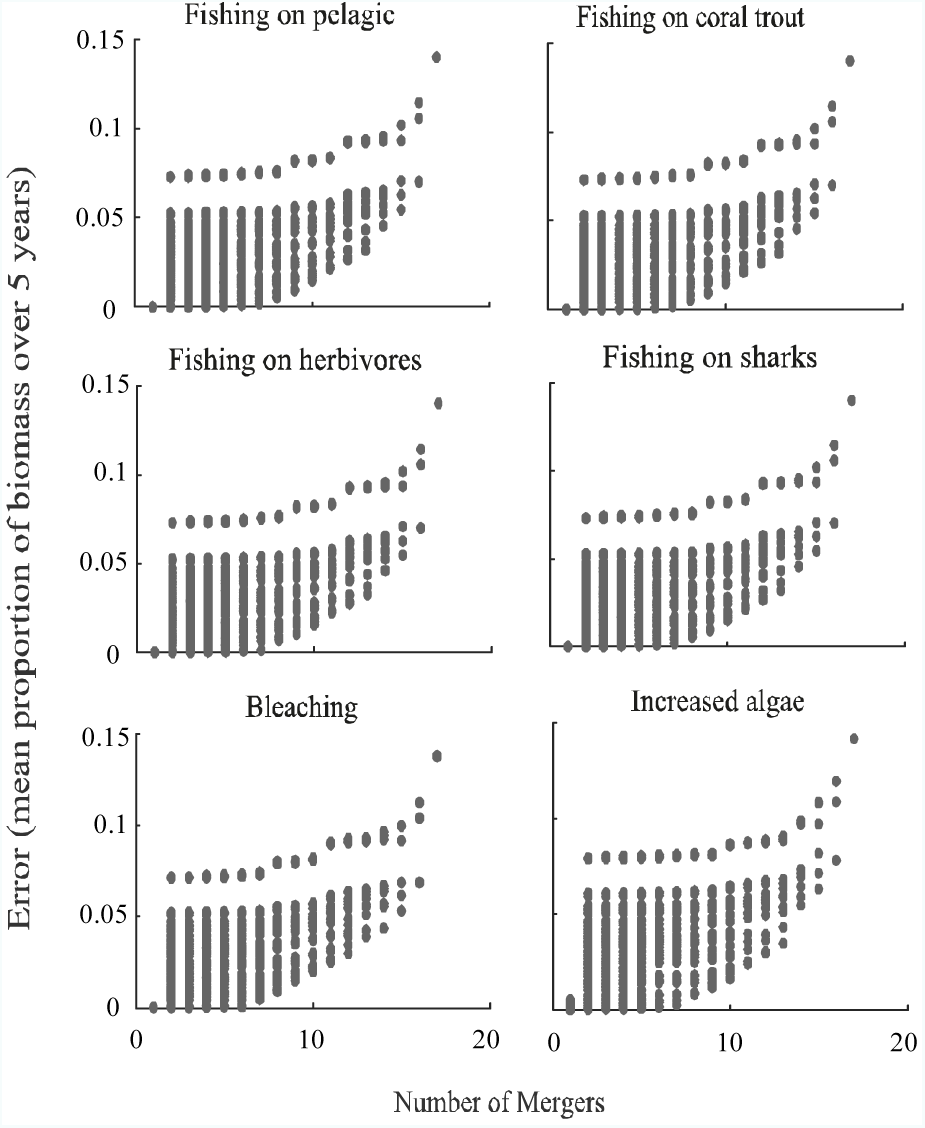
Error estimates for threat scenario 1) to 6) of intensity 0.5 for lutjanids. Each point represents one possible merger of two nodes originating from the model identified with the lowest error in the previous amount of mergers.

**Figure 4.**
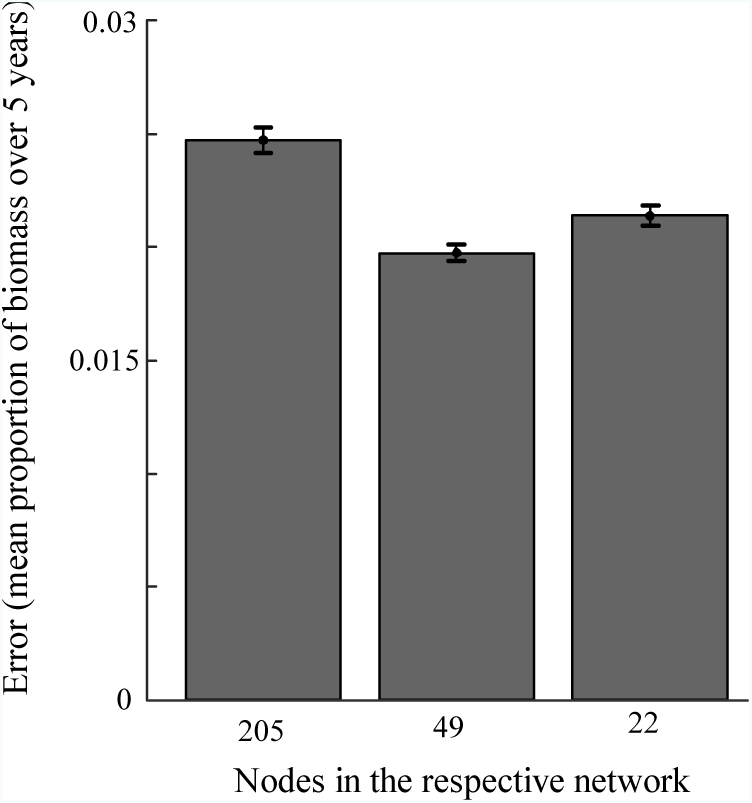
Average error estimates with standard errors for the different resolutions.

### 3.2 Parameter versus structural uncertainty

To compare the balance of parameter and structural uncertainty we are looking at the combined uncertainty (Fig.4). Since 205 nodes is the highest resolution considered here (reference point for the structural uncertainty), it only includes parameter uncertainty with the structural uncertainty being zero (according to our assumption). The 49 and the 22 node model both include structural and parameter uncertainty. We found that the 49 node model was the optimal model in the sense that the overall (structural and parameter) was the lowest. This supports the hypothesis that there is an optimal resolution even when taking into account the parameter uncertainty.

### 3.3 Variable Importance

The bagged trees analysis shows that we can explain most of the variation in the error of a particular merger when considering the how similar in in the predictor variables the two nodes are (*R*^2^=0.89). The variable importance shows that the most effective predictors to determine a good merger are the biomass ratio (∼ node size) and the consumption ratio (∼total amount of inflow into the node) of the species (Fig. 5). Another variable of medium importance is the difference in trophic level between the threat scenario and the merged groups (Fig.5). It is interesting that the similarities in parental or child nodes between nodes to be merged (i.e. similarities in predators and food sources for the merged species) are of low value to determine the direction of merging. In fact, predator overlap is largely associated with all the other predictors (Fig.6), and hence it could be removed from the analysis without any loss of predictive power (*R*^2^ =0.8).

**Figure 5.**
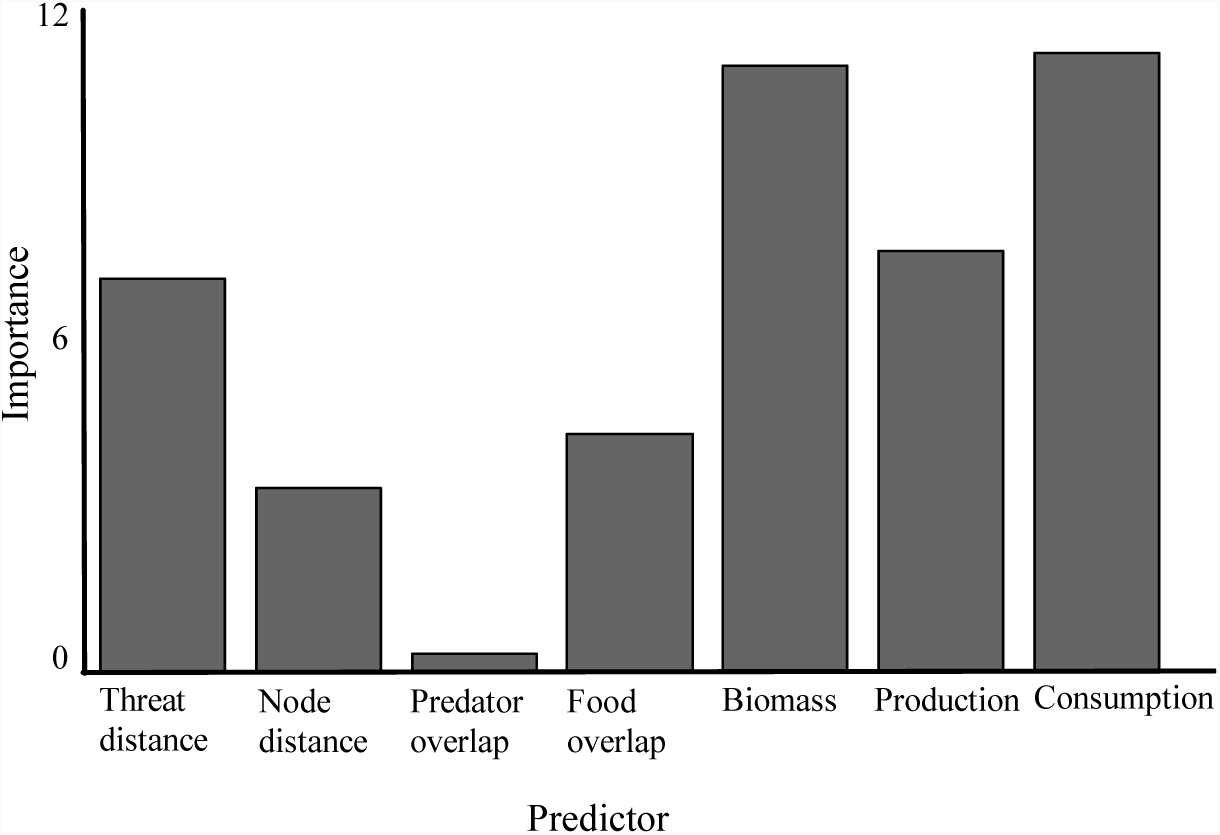
Predictor importance for the regression model (relative measure of importance within the model, no units)

**Figure 6:**
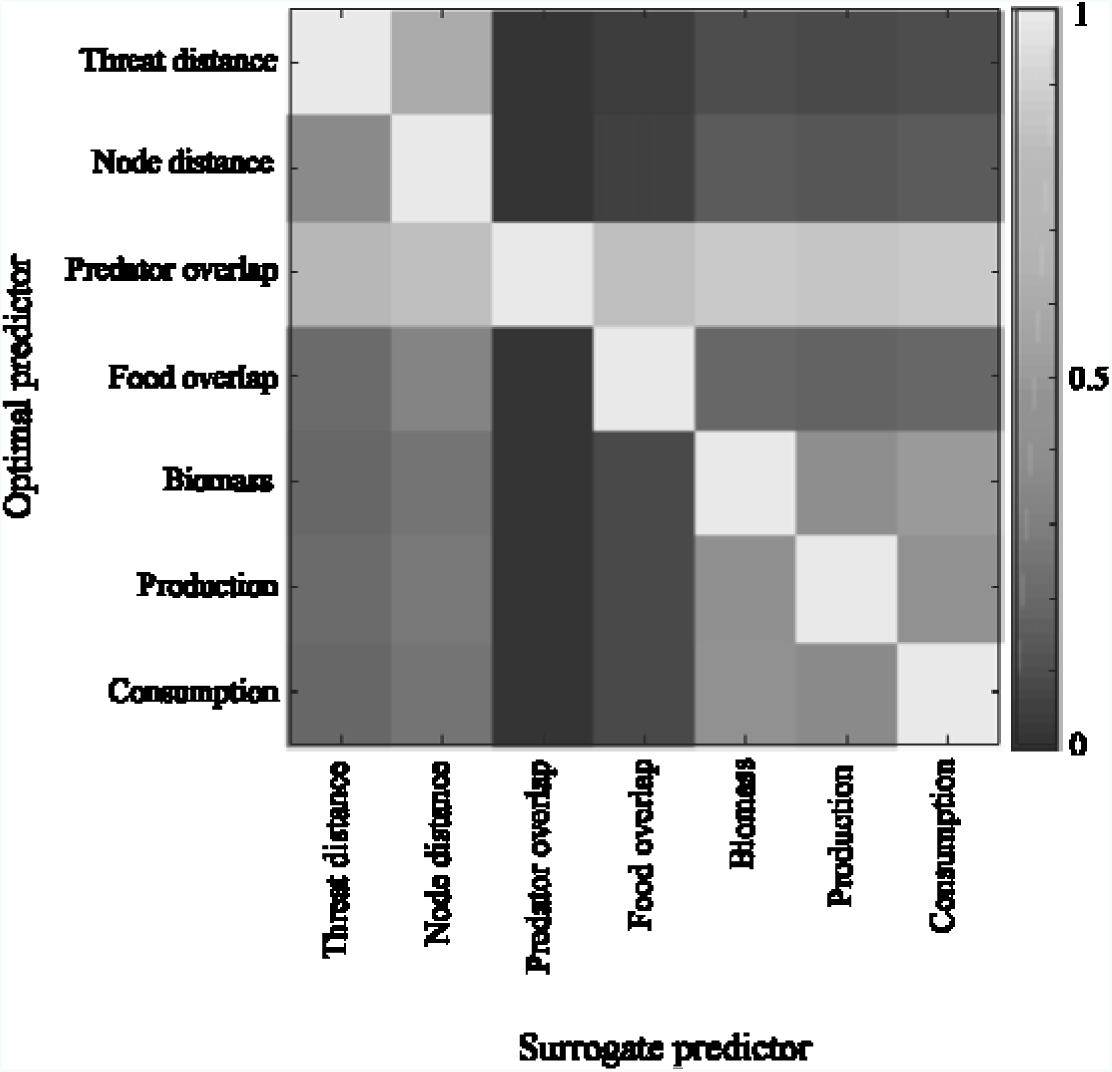
Association between predictors in the tree

## 4 Conclusions and Discussion

This study provides insights into structural uncertainty and more specifically into what role the resolution plays in complex network-based models. Overall, the study gives a good indication of how structural uncertainty in form of resolution could be better integrated into the process of constructing the models based on complex networks. Here we considered an Ecopath model as a prominent example in ecological modelling. We found that different levels of model resolution can change the error in estimating model outcomes in response to exogenous shocks. While higher resolutions always reduce the structural uncertainty this might not be the best resolution overall. Besides the computational capacity needed for high resolution models, the overall (parameter and structural) uncertainty is lowest at a medium resolution. This resolution can be considered as an optimal resolution and can be found by merging species with similar parameters for biomass and total consumption. The study confirms that the distance in trophic level between the merged nodes and the threat can be of importance. Our results highlight that the common practice of putting high importance on the ratios of the size of the merged nodes and their biomasses, rather than on the rest of the network the nodes are connected to, and on the directions of in-and outflows may be warranted.

A result that holds true across all considered functional groups, independently from the number of species or threats is that the minimum error grows with an increasing slope when the model resolution becomes coarser (i.e., with increasing the number of mergers). This means that at a medium resolution level the error is substantially lower than the error at the lowest resolution level (i.e., when a complete merger of all species into one group is achieved). This is not surprising since some species have common ecology, and hence they are even sometimes called “redundant species”, while others differ dramatically in size, food source or predator protection. Redundancy refers to different species full-filling similar ecological roles (Naeem, 1998), hence not much difference can be found if these species are considered as one group in the analysis. While the existence of functional redundancy is still debated (Hoey and Bellwood, 2009), this analysis seems to support the hypothesis. Similar analysis on different systems other than coral reefs might be useful to explore this concept further. The shape of the relationship between the degree of model coarse-graining and the model accuracy suggests to introduce a notion of an optimal grouping of species in terms of structural uncertainty and computational requirements.

This study supports the previous assumption of an opposite hump shape or seesaw between structural and parameter uncertainty (Costanza and Sklar, 1985, Håkanson, 1995, Jester, 1977). When the structural uncertainty is reduced (according to our assumptions entirely removed) and only parameter uncertainty is considered (the 205 node resolution) then we have the highest overall uncertainty. The medium resolution model produced here with 49 nodes shows the lowest overall uncertainty, i.e. while the structural uncertainty is increased the lower amount of parameters resulted in a much lower parameter uncertainty. On the other hand, once the resolution is reduced too much (22 nodes), the structural uncertainty is so high that even combined with the now low parameter uncertainty, the overall uncertainty is increased again. This in conjunction with the shape of the uncertainty estimates across different numbers of mergers supports the hypothesis that an optimal resolution exists. This is not just the optimal across structural uncertainty and computing power required, but also parameter uncertainty. Furthermore, the species grouped in this optimal resolution model (here 49 nodes) are consistent for all of the threat scenarios and intensities. This is especially important when the management questions underlying the model are concerned with multiple threats.

Common practice has often focused on grouping together nodes with similar connections (Cale Jr and Odell, 1980, Fulton, 2001, Gardner and Ashby, 1970, O’Neill, 1975, Wiegert, 1975) rather than the similarity of their abundance in the system. One example is Tudman (2001) who groups all herbivores together irrespective of the large differences in their biomasses ranging from 0.01 to over 10 *t km*^−2^*year*^−1^. On the other hand, it has been recognized that for some very rare species it is better to exclude them from the model than to merge with a species with another group (Fulton, 2001). The results presented here have to be taken with some caution, however, since the data the analysis is based on already assumes some similarity between then species merged, i.e. due to limitations related to computing time the method does not allow a shark and a goby to be merged purely because they have a similar biomass in the system. This restriction here was due to technical reasons mainly, however, it also represents another commonly applied guideline: “do not aggregate serially linked groups” (Fulton et al., 2003), i.e., expert knowledge should be used to provide the initial coarse groupings.

In the literature, it is often suggested that Ecopath models should have as high as possible resolution of the foodweb nodes that are of special interest to the question asked (Heymans et al., 2016, Hollowed et al., 2000, Christensen et al., 2005). For example, if we are investigating fishing, we should differentiate fish groups more explicitly than other parts of the foodweb such as, for example, algae. While this is common practice, it has been pointed out that this method could result in biased results (Fulton et al., 2003). Our study found some support to this guideline, i.e., as we obtained that the difference between the trophic level of the threat and the merged group had some importance, however it needs to be recognised that it was lower than that of the biomass and production. Furthermore, the optimal resolution, and specific species to group in the medium sized model did not change in all of our threat scenarios. Since the threats that were used here can represent anything from bleaching to high trophic level fishing, the results seem to indicate that it is not important which question we are trying to answer when deciding on the species grouped within each node. This contradicts previous advice and should be further investigated, especially, since it is often used as a justification to represent lower trophic levels in massive groups that can represent hundreds or even thousands of species (Tudman, 2001).

The future use of this study is twofold. The study can have a direct use for coral reef models constructed in the future. The optimal groupings found here as well as the total amount of uncertainty found can be utilised for any model of this system. However, the results from this study can reach further since it provides information on how to aggregate nodes in any network model independent on its use. In conclusion, this study is a good foundation for further investigation and the better integration of structural uncertainty in ecosystem, but also other network based models. As long as merging nodes that are not serially linked, the most important determinant of uncertainty is the size ratio of the merged nodes and their total outflow. This can give guidance to future models to manage uncertainty caused by a coarser resolution which modellers have to accept in return for feasible computing resources.

## 5 Acknowledgements

This research has been mostly supported by the YSSP summer fellowship of the International Institute of Applied System Analysis (IIASA). It has also received some support from the HPC at James Cook University and at Queensland University of Technology.

