## Supplementary material for "Optimizing functional groups in ecosystem models: Case study of the Great Barrier Reef": Supplemantary Materials S1-S4

### Supplementary materials

#### *S1: Inputs into the model used*

Table S1.1: Summary table of data inputs into the Ecosim model

| Node | Biomass | Production per biomass | Consumption per biomass | Ecotrophic efficiency | Unassimilated food |
|---|---|---|---|---|---|
| ***sharks/rays*** | 0.2 | 0.2 | 6 | 0.3 | 0.1 |
| ***Plectropomus leopardus*** | 2.91 | 0.38 | 3.7 | 0.48779 | 0.1 |
| ***Plectropomus laevis*** | 0.29 | 0.37 | 4.2 | 0.50097 | 0.1 |
| ***pelagics*** | 1.96 | 0.72 | 8.9 | 0.79893 | 0.1 |
| ***Cephalopholis boenak*** | 0.15 | 1.13 | 10.1 | 0.17306 | 0.1 |
| ***Cephalopholis cyanostigma*** | 0.52 | 0.92 | 9 | 0.21257 | 0.1 |
| ***Cephalopholis mircoprion*** | 0.02 | 1.2 | 11.3 | 0.16296 | 0.1 |
| ***Epinephelus merra*** | 0.03 | 1 | 2.03 | 0.19558 | 0.1 |
| ***Epinephelus ongus*** | 0.01 | 0.92 | 8.3 | 0.21257 | 0.1 |
| ***Aethaloperca rogaa*** | 0.01 | 0.63 | 6.5 | 0.25266 | 0.1 |
| ***Anyperodon leucogrammicus*** | 0.22 | 0.7 | 7 | 0.22736 | 0.1 |
| ***Cephalopholis argus*** | 0.05 | 0.67 | 6.2 | 0.23755 | 0.1 |
| ***Cromileptes altivelis*** | 0.19 | 0.56 | 6 | 0.28421 | 0.1 |
| ***Epinephelus coeruleopunctatus*** | 0.01 | 0.53 | 5.3 | 0.30034 | 0.1 |
| ***Epinephelus cyanopodus*** | 0.01 | 0.38 | 3.6 | 0.41889 | 0.1 |
| ***Epinephelus fasciatus*** | 0.02 | 0.84 | 8.2 | 0.18952 | 0.1 |

| Node | Biomass | Production per biomass | Consumption per biomass | Ecotrophic efficiency | Unassimilated food |
|---|---|---|---|---|---|
| ***Epinephelus fuscoguttatus*** | 0.02 | 0.39 | 4.1 | 0.4082 | 0.1 |
| ***Epinephelus howlandi*** | 0.01 | 0.78 | 7.3 | 0.20407 | 0.1 |
| ***Epinephelus lanceolatus*** | 0.02 | 0.22 | 2.6 | 0.72363 | 0.1 |
| ***Epinephelus macrospilos*** | 0.02 | 0.7 | 6.8 | 0.22743 | 0.1 |
| ***Epinephelus maculatus*** | 0.02 | 0.62 | 6 | 0.25677 | 0.1 |
| ***Epinephelus malabaricus*** | 0.01 | 0.24 | 2.7 | 0.66324 | 0.1 |
| ***Epinephelus polyphekadion*** | 0.04 | 0.47 | 4.5 | 0.33862 | 0.1 |
| ***Lutjanus argentimaculatus*** | 0.04 | 0.36 | 4.2 | 0.54135 | 0.1 |
| ***Lutjaus bohar*** | 0.55 | 0.61 | 5.6 | 0.31946 | 0.1 |
| ***Lutjanus carponotatus*** | 0.16 | 0.2 | 8.3 | 0.97435 | 0.1 |
| ***Lutjanus gibbus*** | 0.04 | 0.78 | 6.9 | 0.24985 | 0.1 |
| ***Lutjanus lemniscatus*** | 0.04 | 0.59 | 6.2 | 0.33031 | 0.1 |
| ***Lutjanus monostigma*** | 0.01 | 0.47 | 6 | 0.41474 | 0.1 |
| ***Lutjanus rivulatus*** | 0.01 | 0.43 | 5.6 | 0.45332 | 0.1 |
| ***Lutjanus russellii*** | 0.43 | 0.71 | 6.7 | 0.27446 | 0.1 |
| ***Lutjanus sebae*** | 0.048 | 0.15 | 3.9 | 0.81201 | 0.1 |
| ***Macalor macularis*** | 0.01 | 0.63 | 6.5 | 0.30941 | 0.1 |
| ***Macalor niger*** | 0.04 | 0.54 | 5.7 | 0.3609 | 0.1 |
| ***Lutjanus flaviflamma*** | 1.46 | 0.69 | 9.1 | 0.31775 | 0.1 |
| ***Lutjanus fulvus*** | 0.01 | 0.84 | 7.5 | 0.26116 | 0.1 |
| ***Lutjanus kasmira*** | 0.01 | 0.83 | 8.9 | 0.26431 | 0.1 |

| Node | Biomass | Production per biomass | Consumption per biomass | Ecotrophic efficiency | Unassimilated food |
|---|---|---|---|---|---|
| ***Lutjanus lutjanus*** | 0.01 | 1.42 | 10.4 | 0.15449 | 0.1 |
| ***Lutjanus quinquelineatus*** | 4.41 | 0.15 | 7.6 | 0.97443 | 0.1 |
| ***Symphorichthys spilurus*** | 0.07 | 0.63 | 6.5 | 0.34801 | 0.1 |
| ***Symphorus nematophorus*** | 0.04 | 0.43 | 4.3 | 0.5099 | 0.1 |
| ***Chaetodon aureofasciatus*** | 0.18 | 1.9 | 16.8 | 0.08443 | 0.1 |
| ***Chaetodon baronessa*** | 0.29 | 1.67 | 15.1 | 0.096059 | 0.1 |
| ***Chaetodon citrinellus*** | 0.02 | 1.85 | 13.7 | 0.086708 | 0.1 |
| ***Chaetodon kleinii*** | 0.02 | 1.67 | 15.1 | 0.096054 | 0.1 |
| ***Chaetodon lineolatus*** | 2.26 | 1.02 | 9.4 | 0.15728 | 0.1 |
| ***Chaetodon melannotus*** | 0.27 | 1.67 | 11.5 | 0.096061 | 0.1 |
| ***Chaetodon plebeius*** | 0.46 | 1.67 | 13.4 | 0.09606 | 0.1 |
| ***Chaetodon rafflesii*** | 0.07 | 1.67 | 15.1 | 0.09606 | 0.1 |
| ***Chaetodon rainfordi*** | 0.51 | 1.67 | 15.1 | 0.096061 | 0.1 |
| ***Chaetodon speculum*** | 0.09 | 1.47 | 13.5 | 0.10913 | 0.1 |
| ***Chaetodon trifascialis*** | 0.17 | 1.47 | 11.1 | 0.10913 | 0.1 |
| ***Chaetodon vagabundus*** | 0.06 | 1.24 | 11.6 | 0.12938 | 0.1 |
| ***Heniochus varius*** | 0.17 | 1.41 | 13 | 0.11378 | 0.1 |
| ***Labrichthys unilineatus*** | 0.24 | 1.5 | 13.7 | 0.10695 | 0.1 |
| ***Arothron nigropunctatus*** | 0.41 | 0.96 | 9.4 | 0.16711 | 0.1 |

| Node | Biomass | Production per biomass | Consumption per biomass | Ecotrophic efficiency | Unassimilated food |
|---|---|---|---|---|---|
| ***Lethrinus miniatus*** | 0.48 | 0.42 | 4.7 | 0.62601 | 0.1 |
| ***Lethrinus atkinsoni*** | 0.16 | 0.6 | 6.4 | 0.25701 | 0.1 |
| ***Lethrinus lentjan*** | 0.01 | 0.53 | 6.1 | 0.291 | 0.1 |
| ***Lethrinus nebulosus*** | 1.84 | 0.43 | 5.6 | 0.35862 | 0.1 |
| ***Lethrinus obsoletus*** | 0.19 | 0.63 | 5.7 | 0.24477 | 0.1 |
| ***Lethrinus olivaceus*** | 0.01 | 0.47 | 5.2 | 0.32815 | 0.1 |
| ***Lethrinus ornatus*** | 0.01 | 1.02 | 8.7 | 0.1512 | 0.1 |
| ***Monotaxis grandoculis*** | 0.63 | 0.63 | 5.5 | 0.24477 | 0.1 |
| ***Myripristis hexagona*** | 0.24 | 1.02 | 8.7 | 0.31515 | 0.1 |
| ***Myripristis kuntee*** | 0.08 | 1.37 | 9.6 | 0.23463 | 0.1 |
| ***Myripristis murdjan*** | 0.62 | 1.02 | 9.9 | 0.31515 | 0.1 |
| ***Myripristis violacea*** | 1.19 | 1.24 | 9.3 | 0.25924 | 0.1 |
| ***Caesio caerulurea*** | 0.6 | 0.92 | 8.3 | 0.34941 | 0.1 |
| ***Pterocaesio digramma*** | 4.93 | 1.02 | 8.4 | 0.31515 | 0.1 |
| ***Caesio cuning*** | 8.61 | 0.63 | 6.5 | 0.51025 | 0.1 |
| ***Cheilodipterus macrodon*** | 0.08 | 1.2 | 9.2 | 0.72113 | 0.1 |
| ***Heteropriacanthus cruentatus*** | 0.12 | 0.71 | 6.4 | 0.45276 | 0.1 |
| ***Aulostomus chinensis*** | 0.216 | 0.51 | 4.3 | 0.8738 | 0.1 |
| ***Balistapus undulatus*** | 0.26 | 1.02 | 9.9 | 0.52404 | 0.1 |
| ***Sufflamen chrysopterum*** | 0.1 | 1.02 | 7.9 | 0.52419 | 0.1 |

| Node | Biomass | Production per biomass | Consumption per biomass | Ecotrophic efficiency | Unassimilated food |
|---|---|---|---|---|---|
| ***Caracanthus unipinna*** | 0.01 | 3.63 | 29.1 | 0.14731 | 0.1 |
| ***Cordion chrysozonus*** | 0.02 | 1.67 | 15.1 | 0.32014 | 0.1 |
| ***Platax teira*** | 0.5 | 0.63 | 14.4 | 0.84854 | 0.1 |
| ***Koumansetta rainfordi*** | 0.01 | 3.01 | 24.9 | 0.29077 | 0.1 |
| ***Asterropteryx semipunctata*** | 0.01 | 3.01 | 24.9 | 0.29077 | 0.1 |
| ***Gobiodon quinquestrigatus*** | 0.01 | 4.67 | 36.8 | 0.18741 | 0.1 |
| ***Gobiodon rivulatus*** | 0.01 | 3.63 | 29.1 | 0.24111 | 0.1 |
| ***Paragobiodon echinocephalus*** | 0.01 | 4.28 | 33.5 | 0.20449 | 0.1 |
| ***Paragobiodon xanthosoma*** | 0.01 | 4.28 | 33.5 | 0.20449 | 0.1 |
| ***Signigobus biocellatus*** | 0.01 | 3.01 | 24.9 | 0.29077 | 0.1 |
| ***Valenciennea strigata*** | 0.75 | 1.47 | 13.5 | 0.031747 | 0.1 |
| ***Plectorhinchus chaetodonoides*** | 0.78 | 0.55 | 5 | 0.81007 | 0.1 |
| ***Plectorhinchus diagrammus*** | 0.28 | 0.84 | 8.3 | 0.63654 | 0.1 |
| ***Plectorhinchus lineatus*** | 0.58 | 0.58 | 5.6 | 0.92179 | 0.1 |
| ***Plectorhinchus picus*** | 0.855 | 0.37 | 5.1 | 0.96324 | 0.1 |
| ***Anampses neoguinaicus*** | 0.042 | 0.4 | 12.7 | 0.95468 | 0.1 |
| ***Bodianus axillaris*** | 0.196 | 0.4 | 12.7 | 0.95483 | 0.1 |
| ***Bodianus mesothorax*** | 0.056 | 0.4 | 11.1 | 0.95451 | 0.1 |
| ***Cheilinus fasciatus*** | 0.574 | 0.4 | 8.3 | 0.95472 | 0.1 |

| Node | Biomass | Production per biomass | Consumption per biomass | Ecotrophic efficiency | Unassimilated food |
|---|---|---|---|---|---|
| ***Cheilinus oxycephalus*** | 0.224 | 0.4 | 14 | 0.95443 | 0.1 |
| ***Cheilinus tribatus*** | 0.966 | 0.4 | 6.7 | 0.95476 | 0.1 |
| ***Choerodon fasciatus*** | 0.196 | 0.4 | 9.9 | 0.95483 | 0.1 |
| ***Cirrhilabrus temminickii*** | 0.126 | 0.4 | 19.3 | 0.95478 | 0.1 |
| ***Coris variegata*** | 0.126 | 0.26 | 12.7 | 0.97938 | 0.1 |
| ***Epibulus insidiator*** | 0.35 | 0.4 | 7 | 0.95491 | 0.1 |
| ***Gomphosus varius*** | 0.364 | 0.4 | 9.9 | 0.95449 | 0.1 |
| ***Halichoeres melanurus*** | 0.052 | 0.21 | 17.2 | 0.97919 | 0.1 |
| ***Hemigymnus fasciatus*** | 0.224 | 0.46 | 5.5 | 0.82994 | 0.1 |
| ***Hemigymnus melapterus*** | 0.196 | 0.4 | 5.1 | 0.95483 | 0.1 |
| ***Labroides dimidiatus*** | 0.09 | 0.12 | 17.8 | 0.99007 | 0.1 |
| ***Labropsis manabei*** | 0.014 | 0.4 | 19.2 | 0.95488 | 0.1 |
| ***Novaculichthys taeniorus*** | 0.028 | 0.4 | 9.9 | 0.95471 | 0.1 |
| ***Oxycheilinus dgramma*** | 0.854 | 0.4 | 8.3 | 0.95468 | 0.1 |
| ***Pseudocheilinus evanidus*** | 0.014 | 0.4 | 21.9 | 0.95488 | 0.1 |
| ***Pseudocheilinus hexataenia*** | 0.042 | 0.4 | 19.2 | 0.95468 | 0.1 |
| ***Thalassoma amlycephalum*** | 0.014 | 0.4 | 14.5 | 0.95488 | 0.1 |
| ***Thalassoma hardwicke*** | 0.112 | 0.4 | 12.6 | 0.95482 | 0.1 |
| ***Thalassoma lunare*** | 0.07 | 0.4 | 10.5 | 0.95456 | 0.1 |
| ***Wetmorella nigropinnata*** | 0.014 | 0.4 | 21.9 | 0.95488 | 0.1 |

| **Node** | **Biomass** | **Production per biomass** | **Consumption per biomass** | **Ecotrophic efficiency** | **Unassimilated food** |
|---|---|---|---|---|---|
| ***Cantherhines sandwichiensis*** | 0.1 | 1.4 | 28.5 | 0.4488 | 0.1 |
| ***Paraluteres prionurus*** | 0.01 | 2.08 | 40 | 0.3021 | 0.1 |
| ***Pervagor melanocephalus*** | 0.09 | 1.6 | 31.9 | 0.3927 | 0.1 |
| ***Scolopsis bilineata*** | 1.03 | 1.24 | 10.1 | 0.43117 | 0.1 |
| ***Ostracion cubicus*** | 0.12 | 0.77 | 14.6 | 0.69481 | 0.1 |
| ***Ostracion meleagris*** | 0.01 | 1.17 | 24.2 | 0.45703 | 0.1 |
| ***Pempheris oualensis*** | 0.42 | 1.37 | 12.7 | 0.39029 | 0.1 |
| ***Assesor macneilli*** | 0.03 | 3.18 | 25.2 | 0.16812 | 0.1 |
| ***Plesiops coeruleolineatus*** | 0.04 | 2.22 | 19.2 | 0.24078 | 0.1 |
| ***Centropyge bicolor*** | 0.15 | 1.67 | 15.1 | 0.32027 | 0.1 |
| ***Centropyge bispinosa*** | 0.04 | 2.22 | 15.9 | 0.24078 | 0.1 |
| ***Centropyge vrolikii*** | 0.22 | 2.22 | 19.2 | 0.2409 | 0.1 |
| ***Chaetodontoplus personifer*** | 0.1 | 0.92 | 19.9 | 0.58117 | 0.1 |
| ***Pomacanthus sexstriatus*** | 0.5 | 0.76 | 16.9 | 0.7034 | 0.1 |
| ***Pygoplites diacanthus*** | 0.1 | 1.17 | 24.4 | 0.45699 | 0.1 |
| ***Pseudochromis flammicauda*** | 0.01 | 3.41 | 27.5 | 0.15681 | 0.1 |
| ***Pseudochromis fuscus*** | 0.01 | 2.4 | 20.5 | 0.2228 | 0.1 |
| ***Pterois antennata*** | 0.07 | 1.37 | 12.7 | 0.39031 | 0.1 |
| ***Sphyraena pinguis*** | 0.36 | 1.02 | 9.9 | 0.52428 | 0.1 |
| ***Zanclus cornutus*** | 0.19 | 1.24 | 9.2 | 0.43101 | 0.1 |
| ***Nectamia bandanensis*** | 0.1 | 2.22 | 16.2 | 0.44451 | 0.1 |

| Node | Biomass | Production per biomass | Consumption per biomass | Ecotrophic efficiency | Unassimilated food |
|---|---|---|---|---|---|
| ***Apogon coccineus*** | 0.01 | 3.18 | 26.1 | 0.31034 | 0.1 |
| ***Ostorhinchus cyanosoma*** | 0.01 | 5.01 | 18.8 | 0.19698 | 0.1 |
| ***Apogonichthyoides taeniatus*** | 0.01 | 1.53 | 14 | 0.64502 | 0.1 |
| ***Pristicon trimaculatus*** | 0.25 | 1.6 | 13.7 | 0.61677 | 0.1 |
| ***Cheilodipterus quinquelineatus*** | 0.02 | 1.85 | 15.6 | 0.53349 | 0.1 |
| ***Fowleria punctulata*** | 0.01 | 2.5 | 21.2 | 0.39475 | 0.1 |
| ***Sargocentron diadema*** | 0.35 | 2.13 | 13.3 | 0.20799 | 0.1 |
| ***Acanthochromis polyacanthus*** | 1.94 | 2.08 | 18.2 | 0.21295 | 0.1 |
| ***Amblyglyphidodon aureus*** | 0.01 | 1.95 | 17.2 | 0.22717 | 0.1 |
| ***Amblyglyphidodon curacao*** | 3.56 | 2.4 | 16.1 | 0.18456 | 0.1 |
| ***Amblyglyphidodon leucogaster*** | 0.41 | 1.85 | 16.4 | 0.2394 | 0.1 |
| ***Amphiprion akindynos*** | 0.04 | 2.4 | 16.4 | 0.1846 | 0.1 |
| ***Amphiprion clarkii*** | 0.02 | 1.67 | 15.1 | 0.2653 | 0.1 |
| ***Amphiprion melanopus*** | 0.02 | 1.42 | 21.2 | 0.31201 | 0.1 |
| ***Chromis aboinensis*** | 0.01 | 2.4 | 20.5 | 0.18457 | 0.1 |
| ***Chromis atripectoralis*** | 8.33 | 1.95 | 13.4 | 0.22715 | 0.1 |
| ***Chromis atripes*** | 0.18 | 2.4 | 20.5 | 0.18457 | 0.1 |
| ***Chromis lepidolepis*** | 0.01 | 2.6 | 21.9 | 0.17038 | 0.1 |
| ***Chromis margaritifer*** | 0.01 | 2.4 | 20.5 | 0.18457 | 0.1 |

| Node | Biomass | Production per biomass | Consumption per biomass | Ecotrophic efficiency | Unassimilated food |
|---|---|---|---|---|---|
| ***Chromis nitida*** | 0.01 | 2.87 | 23.9 | 0.15435 | 0.1 |
| ***Chromis retrofasciata*** | 0.01 | 4.28 | 36.8 | 0.1035 | 0.1 |
| ***Chromis ternatensis*** | 0.88 | 6.61 | 17.8 | 0.067008 | 0.1 |
| ***Chromis weberi*** | 0.05 | 1.85 | 16.4 | 0.23942 | 0.1 |
| ***Chromis xanthura*** | 0.04 | 1.67 | 15.1 | 0.2653 | 0.1 |
| ***Chrysptera rex*** | 0.01 | 2.87 | 23.9 | 0.15435 | 0.1 |
| ***Chrysptera rollandi*** | 0.04 | 4.28 | 33.5 | 0.10352 | 0.1 |
| ***Chrysptera talboti*** | 0.01 | 3.18 | 26.1 | 0.1393 | 0.1 |
| ***Neopomacentrus azysron*** | 1.93 | 3.18 | 21.6 | 0.13929 | 0.1 |
| ***Pomacentrus brachialis*** | 0.31 | 2.6 | 18.1 | 0.24007 | 0.1 |
| ***Pomacentrus coelestis*** | 0.02 | 2.4 | 18.9 | 0.26008 | 0.1 |
| ***Pomacentrus lepidogenys*** | 1.22 | 2.4 | 15.9 | 0.26003 | 0.1 |
| ***Pomacentrus melanochir*** | 0.01 | 2.87 | 23.9 | 0.21747 | 0.1 |
| ***Pomacentrus moluccensis*** | 0.93 | 1.67 | 24.2 | 0.37371 | 0.1 |
| ***Pomacentrus philippinus*** | 0.22 | 2.22 | 15.9 | 0.19956 | 0.1 |
| ***Acanthurus mata*** | 1.25 | 0.71 | 16.1 | 0.83139 | 0.1 |
| ***Ctenochaetus striatus*** | 0.9 | 1.6 | 21.2 | 0.26779 | 0.1 |
| ***Ctenochaetus binotatus*** | 0.24 | 1.28 | 25.2 | 0.33474 | 0.1 |
| ***Acanthurus lineatus*** | 0.31 | 0.44 | 24.3 | 0.96282 | 0.1 |
| ***Acanthurus nigrofuscus*** | 0.286 | 0.2 | 37.9 | 0.96283 | 0.1 |

| Node | Biomass | Production per biomass | Consumption per biomass | Ecotrophic efficiency | Unassimilated food |
|---|---|---|---|---|---|
| ***Acanthurus triostegus*** | 0.13 | 0.81 | 32.2 | 0.52302 | 0.1 |
| ***Nasa unicornis*** | 0.75 | 0.32 | 19.7 | 0.81821 | 0.1 |
| ***Zebrasoma scopas*** | 0.56 | 1.37 | 34.4 | 0.19111 | 0.1 |
| ***Cetoscraus bicolor*** | 0.06 | 0.47 | 17.4 | 0.55707 | 0.1 |
| ***Chlorurus gibbus*** | 1.66 | 0.51 | 23.3 | 0.51339 | 0.1 |
| ***Chlorurus sordidus*** | 2.324 | 0.19 | 33.7 | 0.98431 | 0.1 |
| ***Chlorurus bleekeri*** | 0.01 | 0.72 | 25.1 | 0.36366 | 0.1 |
| ***Scarus prasiognathos*** | 1.58 | 0.56 | 20.3 | 0.46755 | 0.1 |
| ***Scarus dubius*** | 0.36 | 0.91 | 30.5 | 0.28772 | 0.1 |
| ***Scarus festivus*** | 0.6 | 0.77 | 26.4 | 0.34004 | 0.1 |
| ***Scarus flavipectoralis*** | 0.09 | 1.02 | 33.7 | 0.2567 | 0.1 |
| ***Scarus forsteni*** | 1.15 | 0.67 | 23.4 | 0.39079 | 0.1 |
| ***Scarus frenatus*** | 10.032 | 0.12 | 19.3 | 0.99177 | 0.1 |
| ***Scarus globiceps*** | 1.45 | 1.1 | 35.9 | 0.23803 | 0.1 |
| ***Scarus niger*** | 2.2 | 0.84 | 24.5 | 0.3117 | 0.1 |
| ***Scarus psittacus*** | 0.27 | 0.79 | 20.7 | 0.33143 | 0.1 |
| ***Scarus spinus*** | 0.01 | 1.02 | 33.7 | 0.2567 | 0.1 |
| ***Siganus corallinus*** | 1.07 | 1.08 | 44 | 0.24243 | 0.1 |
| ***Siganus doliatus*** | 0.3 | 1.2 | 31.4 | 0.21819 | 0.1 |
| ***Siganus punctatus*** | 0.6 | 0.84 | 28.4 | 0.3117 | 0.1 |
| ***Siganus vulpinus*** | 0.32 | 1.2 | 38.6 | 0.21819 | 0.1 |
| ***small hebivores*** | 17 | 0.6 | 17.5 | 0.58769 | 0.1 |
| ***cephalopods*** | 1.88 | 4.59 | 17.5 | 0.76562 | 0.1 |
| ***large invertebrates*** | 80 | 1.64 | 15 | 0.97045 | 0.1 |
| ***small invertebrates*** | 70 | 3.08 | 11.8 | 0.92594 | 0.1 |
| ***zooplankton*** | 60 | 55 | 165 | 0.93757 | 0.4 |
| ***sponges*** | 44 | 0.96 | 3 | 0.94951 | 0.1 |

| Node | Biomass | Production per biomass | Consumption per biomass | Ecotrophic efficiency | Unassimilated food |
|---|---|---|---|---|---|
| ***coral/zooxanthallae*** | 150 | 1.1 | 7.3 | 0.60813 | 0.1 |
| ***phytoplankton*** | 12.8 | 475 | 0 | 0.66703 | 0 |
| ***epiphilic algae*** | 100 | 25 | 0 | 0.3772 | 0 |
| ***Detritus*** | 135 | 0 | 0 | 0.75506 | 0 |

***S2: Supplementary methods***

Consider equation (3)

$$Pb_i B_i = \sum_j Qb_j B_j DC_{ij} + E_i + Pb_i B_i (1 - EE_i) \quad \text{(S2-1)}$$

First we denote the nodes to be merged as $n$ and $n-1$ and the newly created node as $n-1$ in the new system. This does not reduce generality since all nodes can be renumbered to be included into the merger. The parameters of the new system will be denoted as $\hat{B}, \widehat{Pb}, \widehat{Qb}, \widehat{EE}, \hat{E}, \widehat{DC}$. Since nodes $1, \dots, n-2$ are not being merged, it can be assumed that

$$\hat{E}_i = E_i \quad \text{(S2-2)}$$

$$\widehat{Qb}_i = Qb_i \quad \text{(S2-3)}$$

$$\widehat{Pb}_i = Pb_i \quad \text{(S2-4)}$$

$$\widehat{EE}_i = EE_i \text{ for all } i = 1, \dots, n-2 \quad \text{(S2-5)}$$

and

$$\widehat{DC}_{ij} = DC_{ij} \text{ for all } i, j = 1, \dots, n-2. \quad \text{(S2-6)}$$

Since the equation (1-3) is linear

$$\hat{B}_{n-1} = B_{n-1} + B_n \quad \text{(S2-7)}$$

and

$$\hat{E}_{n-1} = E_{n-1} + E_n \quad \text{(S2-8)}$$

To fit the n-1 dimensional system we need to define the parameters for $\widehat{Qb}_{n-1}, \widehat{Pb}_{n-1}, \widehat{EE}_{n-1}$, $\widehat{DC}_{n-1,j}$ and $\widehat{DC}_{i,n-1}$ for all $i, j = 1, \dots, n-1$. Since the total consumption and production are additive properties, we can calculate the $\widehat{Qb}_{n-1}$ and $\widehat{Pb}_{n-1}$ as weighted averages.

$$\widehat{Qb}_{n-1} = \frac{\hat{Q}_{n-1}}{\hat{B}_{n-1}} = \frac{Q_{n-1} + Q_n}{B_{n-1} + B_n} \quad \text{(S2-9)}$$

$$\widehat{Pb}_{n-1} = \frac{\hat{P}_{n-1}}{\hat{B}_{n-1}} = \frac{P_{n-1} + P_n}{B_{n-1} + B_n} \quad \text{(S2-10)}$$

Furthermore the energy ‘lost’, not passed onto the next trophic level is additive.

$$\widehat{EE}_{n-1} \hat{P}_{n-1} = EE_{n-1} P_{n-1} + EE_n P_n \quad \text{(S2-11)}$$

Since the steady state productions are known we can calculate

$$\widehat{EE}_{n-1} = \frac{EE_{n-1} P_{n-1} + EE_n P_n}{P_{n-1} + P_n} \quad \text{(S2-12)}$$

The proportion of the consumption of a predator $j$ is also an additive property resulting in

$$\widehat{DC}_{n-1,j} = DC_{nj} + DC_{n-1,j} \quad \text{(S2-13)}$$

Furthermore, the total part of the diet of a predator $j$ made up of prey $i$ is additive.

$$\widehat{DC}_{n-1,j} = \frac{DC_{nj}Q_n + DC_{n-1,j}Q_{n-1}}{Q_n + Q_{n-1}} \quad \text{(S2-14)}$$

Let's calculate the $\hat{B}_i$ for $i = 1, \ldots, n-2$ using (1-3) and substituting in (3-1) to (3-3).

$$\hat{B}_i = \frac{\sum_j \widehat{Qb}_j \hat{B}_j \widehat{DC}_{ji} + \hat{E}_i}{\widehat{EE}_i \widehat{Pb}_i} \quad \text{(S2-15)}$$

$$\hat{B}_i = \frac{\sum_j^{n-2} Qb_j \hat{B}_j DC_{ji} + \widehat{Qb}_{n-1} \hat{B}_{n-1} \widehat{DC}_{n-1i} + E_i}{EE_i Pb_i} \quad \text{(S2-16)}$$

Substitute in (S2-63), (S2-9), (S2-10) and (S2-14)

$$\hat{B}_i = \frac{\sum_j^{n-2} Qb_j \hat{B}_j DC_{ji} + \left(\frac{Q_{n-1} + Q_n}{B_{n-1} + B_n}\right)(B_{n-1} + B_n)\left(\frac{DC_{nj}Q_n + DC_{n-1,j}Q_{n-1}}{Q_n + Q_{n-1}}\right) + E_i}{EE_i Pb_i} \quad \text{(S2-17)}$$

$$\hat{B}_i = \frac{\sum_j^{n-2} Qb_j \hat{B}_j DC_{ji} + DC_{nj}Q_n + DC_{n-1,j}Q_{n-1} + E_i}{EE_i Pb_i} \quad \text{(S2-18)}$$

This means that due to the linearity of the Ecopath equations a system with a reduced number of nodes can be recreated 100% without creating an error.

$$\hat{B}_i = \frac{\sum_j Qb_j B_j DC_{ji} + E_i}{EE_i Pb_i} = B_i \quad \text{(S2-19)}$$

*S3: Species aggregation according to different model resolutions*

**Table S3.1:** Node classification of the species included in this model. The full model refers to the maximum resolution considered here with 206 nodes. Reduced 1 refers to the lowest resolution model similar to Tudman (2001) with 22 nodes. Reduced 2 refers to the model created by evaluating the errors for the individual functional groups adding up to 22 nodes. Reduced 3 includes the nodes found when merging all fish species and includes a total of 34 nodes.

| **Full model** | **Reduced 1** | **Reduced 2** | **Reduced 3** |
|---|---|---|---|
| **sharks/rays** | sharks/rays | sharks/rays | sharks/rays |
| ***Plectropomus leopardus*** | coral trout | coral trout | coral trout |
| ***Plectropomus laevis*** | coral trout | coral trout | coral trout |
| **pelagics** | pelagics | Pelagics | pelagics |
| ***Cephalopholis boenak*** | serranids | serranidsA | FishA |
| ***Cephalopholis cyanostigma*** | serranids | serranidsA | FishA |
| ***Cephalopholis mircoprion*** | serranids | serranidsB | FishB |
| ***Epinephelus merra*** | serranids | serranidsB | FishA |
| ***Epinephelus ongus*** | serranids | serranidsB | FishA |
| ***Aethaloperca rogaa*** | serranids | serranidsB | FishA |
| ***Anyperodon leucogrammicus*** | serranids | serranidsB | FishB |
| ***Cephalopholis argus*** | serranids | serranidsB | FishC |
| ***Cromileptes altivelis*** | serranids | serranidsB | FishC |
| ***Epinephelus coeruleopunctatus*** | serranids | serranidsB | FishD |
| ***Epinephelus cyanopodus*** | serranids | serranidsB | FishA |
| ***Epinephelus fasciatus*** | serranids | serranidsB | FishC |
| ***Epinephelus fuscoguttatus*** | serranids | serranidsB | FishD |
| ***Epinephelus howlandi*** | serranids | serranidsB | FishD |

| Full model | Reduced 1 | Reduced 2 | Reduced 3 |
|---|---|---|---|
| ***Epinephelus lanceolatus*** | serranids | serranidsB | FishC |
| ***Epinephelus macrospilos*** | serranids | serranidsB | FishB |
| ***Epinephelus maculatus*** | serranids | serranidsB | FishE |
| ***Epinephelus malabaricus*** | serranids | serranidsB | FishD |
| ***Epinephelus polyphekadion*** | serranids | serranidsB | FishC |
| ***Lutjanus argentimaculatus*** | lutjanids | lutjanidsA | FishB |
| ***Lutjaus bohar*** | lutjanids | lutjanidsB | FishF |
| ***Lutjanus carponotatus*** | lutjanids | lutjanidsB | FishG |
| ***Lutjanus gibbus*** | lutjanids | lutjanidsB | FishF |
| ***Lutjanus lemniscatus*** | lutjanids | lutjanidsB | FishG |
| ***Lutjanus monostigma*** | lutjanids | lutjanidsA | FishH |
| ***Lutjanus rivulatus*** | lutjanids | lutjanidsB | FishI |
| ***Lutjanus russellii*** | lutjanids | lutjanidsB | FishJ |
| ***Lutjanus sebae*** | lutjanids | lutjanidsA | FishF |
| ***Macalor macularis*** | lutjanids | lutjanidsB | FishJ |
| ***Macalor niger*** | lutjanids | lutjanidsB | FishF |
| ***Lutjanus flaviflamma*** | lutjanids | lutjanidsB | FishA |
| ***Lutjanus fulvus*** | lutjanids | lutjanidsB | FishE |
| ***Lutjanus kasmira*** | lutjanids | lutjanidsB | FishB |
| ***Lutjanus lutjanus*** | lutjanids | lutjanidsB | FishE |
| ***Lutjanus quinquelineatus*** | lutjanids | lutjanidsA | FishA |
| ***Symphorichthys spilurus*** | lutjanids | lutjanidsB | FishI |

| **Full model** | **Reduced 1** | **Reduced 2** | **Reduced 3** |
|---|---|---|---|
| ***Symphorus nematophorus*** | lutjanids | lutjanidsB | FishG |
| ***Chaetodon aureofasciatus*** | corallivores | corallivores | FishK |
| ***Chaetodon baronessa*** | corallivores | corallivores | FishL |
| ***Chaetodon citrinellus*** | corallivores | corallivores | FishM |
| ***Chaetodon kleinii*** | corallivores | corallivores | FishE |
| ***Chaetodon lineolatus*** | corallivores | corallivores | FishI |
| ***Chaetodon melannotus*** | corallivores | corallivores | FishM |
| ***Chaetodon plebeius*** | corallivores | corallivores | FishC |
| ***Chaetodon rafflesii*** | corallivores | corallivores | FishK |
| ***Chaetodon rainfordi*** | corallivores | corallivores | FishF |
| ***Chaetodon speculum*** | corallivores | corallivores | FishM |
| ***Chaetodon trifascialis*** | corallivores | corallivores | FishL |
| ***Chaetodon vagabundus*** | corallivores | corallivores | FishN |
| ***Heniochus varius*** | corallivores | corallivores | FishI |
| ***Labrichthys unilineatus*** | corallivores | corallivores | FishG |
| ***Arothron nigropunctatus*** | corallivores | corallivores | FishN |
| ***Lethrinus miniatus*** | lethrinids | Lethrinids | FishO |
| ***Lethrinus atkinsoni*** | lethrinids | Lethrinids | FishN |
| ***Lethrinus lentjan*** | lethrinids | Lethrinids | FishH |
| ***Lethrinus nebulosus*** | lethrinids | Lethrinids | FishB |
| ***Lethrinus obsoletus*** | lethrinids | Lethrinids | FishG |
| ***Lethrinus olivaceus*** | lethrinids | Lethrinids | FishH |
| ***Lethrinus ornatus*** | lethrinids | Lethrinids | FishF |

| **Full model** | **Reduced 1** | **Reduced 2** | **Reduced 3** |
|---|---|---|---|
| ***Monotaxis grandoculis*** | lethrinids | Lethrinids | FishN |
| ***Myripristis hexagona*** | large planktivores | large planktivores | FishM |
| ***Myripristis kuntee*** | large planktivores | large planktivores | FishK |
| ***Myripristis murdjan*** | large planktivores | large planktivores | FishO |
| ***Myripristis violacea*** | large planktivores | large planktivores | FishG |
| ***Caesio caerulurea*** | large planktivores | large planktivores | FishD |
| ***Pterocaesio digramma*** | large planktivores | large planktivores | FishE |
| ***Caesio cuning*** | large planktivores | large planktivores | FishE |
| ***Cheilodipterus macrodon*** | large planktivores | large planktivores | FishL |
| ***Heteropriacanthus cruentatus*** | large planktivores | large planktivores | FishL |
| ***Aulostomus chinensis*** | other demersals | other_demersalsA | FishA |
| ***Balistapus undulatus*** | other demersals | other_demersalsA | FishP |
| ***Sufflamen chrysopterum*** | other demersals | other_demersalsB | FishM |
| ***Caracanthus unipinna*** | other demersals | other_demersalsC | FishI |
| ***Cordion chrysozonus*** | other demersals | other_demersalsD | FishL |
| ***Platax teira*** | other demersals | other_demersalsC | FishG |
| ***Koumansetta rainfordi*** | other demersals | other_demersalsE | FishM |
| ***Asterropteryx semipunctata*** | other demersals | other_demersalsB | FishL |
| ***Gobiodon quinquestrigatus*** | other demersals | other_demersalsF | FishK |
| ***Gobiodon rivulatus*** | other demersals | other_demersalsD | FishQ |
| ***Paragobiodon echinocephalus*** | other demersals | other_demersalsA | FishN |

| Full model | Reduced 1 | Reduced 2 | Reduced 3 |
|---|---|---|---|
| ***Paragobiodon xanthosoma*** | other demersals | other_demersalsG | FishG |
| ***Signigobus biocellatus*** | other demersals | other_demersalsF | FishK |
| ***Valenciennea strigata*** | other demersals | other_demersalsF | FishR |
| ***Plectorhinchus chaetodonoides*** | other demersals | other_demersalsG | FishO |
| ***Plectorhinchus diagrammus*** | other demersals | other_demersalsG | FishS |
| ***Plectorhinchus lineatus*** | other demersals | other_demersalsG | FishQ |
| ***Plectorhinchus picus*** | other demersals | other_demersalsD | FishI |
| ***Anampses neoguinaicus*** | other demersals | other_demersalsG | FishI |
| ***Bodianus axillaris*** | other demersals | other_demersalsH | FishP |
| ***Bodianus mesothorax*** | other demersals | other_demersalsI | FishG |
| ***Cheilinus fasciatus*** | other demersals | other_demersalsE | FishQ |
| ***Cheilinus oxycephalus*** | other demersals | other_demersalsH | FishG |
| ***Cheilinus tribatus*** | other demersals | other_demersalsD | FishN |
| ***Choerodon fasciatus*** | other demersals | other_demersalsG | FishQ |
| ***Cirrhilabrus temminickii*** | other demersals | other_demersalsJ | FishQ |
| ***Coris variegata*** | other demersals | other_demersalsJ | FishG |
| ***Epibulus insidiator*** | other demersals | other_demersalsJ | FishT |
| ***Gomphosus varius*** | other demersals | other_demersalsI | FishM |
| ***Halichoeres melanurus*** | other demersals | other_demersalsK | FishG |
| ***Hemigymnus fasciatus*** | other demersals | other_demersalsG | FishR |
| ***Hemigymnus melapterus*** | other demersals | other_demersalsG | FishG |

| **Full model** | **Reduced 1** | **Reduced 2** | **Reduced 3** |
|---|---|---|---|
| ***Labroides dimidiatus*** | other demersals | other_demersalsK | FishI |
| ***Labropsis manabei*** | other demersals | other_demersalsG | FishM |
| ***Novaculichthys taeniorus*** | other demersals | other_demersalsA | FishG |
| ***Oxycheilinus dgramma*** | other demersals | other_demersalsB | FishG |
| ***Pseudocheilinus evanidus*** | other demersals | other_demersalsG | FishI |
| ***Pseudocheilinus hexataenia*** | other demersals | other_demersalsL | FishI |
| ***Thalassoma amlycephalum*** | other demersals | other_demersalsA | FishI |
| ***Thalassoma hardwicke*** | other demersals | other_demersalsM | FishO |
| ***Thalassoma lunare*** | other demersals | other_demersalsN | FishQ |
| ***Wetmorella nigropinnata*** | other demersals | other_demersalsL | FishI |
| ***Cantherhines sandwichiensis*** | other demersals | other_demersalsN | FishN |
| ***Paraluteres prionurus*** | other demersals | other_demersalsO | FishG |
| ***Pervagor melanocephalus*** | other demersals | other_demersalsO | FishM |
| ***Scolopsis bilineata*** | other demersals | other_demersalsA | FishC |
| ***Ostracion cubicus*** | other demersals | other_demersalsP | FishG |
| ***Ostracion meleagris*** | other demersals | other_demersalsG | FishG |
| ***Pempheris oualensis*** | other demersals | other_demersalsG | FishG |
| ***Assesor macneilli*** | other demersals | other_demersalsM | FishO |
| ***Plesiops coeruleolineatus*** | other demersals | other_demersalsQ | FishS |
| ***Centropyge bicolor*** | other demersals | other_demersalsQ | FishR |
| ***Centropyge bispinosa*** | other demersals | other_demersalsR | FishS |
| ***Centropyge vrolikii*** | other demersals | other_demersalsL | FishS |

| Full model | Reduced 1 | Reduced 2 | Reduced 3 |
|---|---|---|---|
| ***Chaetodontoplus personifer*** | other demersals | other_demersalsS | FishI |
| ***Pomacanthus sexstriatus*** | other demersals | other_demersalsG | FishT |
| ***Pygoplites diacanthus*** | other demersals | other_demersalsR | FishG |
| ***Pseudochromis flammicauda*** | other demersals | other_demersalsM | FishI |
| ***Pseudochromis fuscus*** | other demersals | other_demersalsM | FishC |
| ***Pterois antennata*** | other demersals | other_demersalsT | FishS |
| ***Sphyraena pinguis*** | other demersals | other_demersalsM | FishC |
| ***Zanclus cornutus*** | other demersals | other_demersalsT | FishG |
| ***Nectamia bandanensis*** | small planktivores | small_planktivoresA | FishG |
| ***Apogon coccineus*** | small planktivores | small_planktivoresA | FishO |
| ***Ostorhinchus cyanosoma*** | small planktivores | small_planktivoresB | FishO |
| ***Apogonichthyoides taeniatus*** | small planktivores | small_planktivoresA | FishN |
| ***Pristicon trimaculatus*** | small planktivores | small_planktivoresA | FishB |
| ***Cheilodipterus quinquelineatus*** | small planktivores | small_planktivoresA | FishS |
| ***Fowleria punctulata*** | small planktivores | small_planktivoresA | FishP |
| ***Sargocentron diadema*** | small planktivores | small_planktivoresA | FishF |
| ***Acanthochromis polyacanthus*** | small planktivores | small_planktivoresA | FishI |
| ***Amblyglyphidodon aureus*** | small planktivores | small_planktivoresA | FishE |
| ***Amblyglyphidodon curacao*** | small planktivores | small_planktivoresB | FishB |
| ***Amblyglyphidodon leucogaster*** | small planktivores | small_planktivoresA | FishI |
| ***Amphiprion akindynos*** | small planktivores | small_planktivoresA | FishG |

| **Full model** | **Reduced 1** | **Reduced 2** | **Reduced 3** |
|---|---|---|---|
| ***Amphiprion clarkii*** | small planktivores | small_planktivoresA | FishR |
| ***Amphiprion melanopus*** | small planktivores | small_planktivoresA | FishI |
| ***Chromis aboinensis*** | small planktivores | small_planktivoresA | FishM |
| ***Chromis atripectoralis*** | small planktivores | small_planktivoresA | FishC |
| ***Chromis atripes*** | small planktivores | small_planktivoresC | FishG |
| ***Chromis lepidolepis*** | small planktivores | small_planktivoresC | FishK |
| ***Chromis margaritifer*** | small planktivores | small_planktivoresA | FishL |
| ***Chromis nitida*** | small planktivores | small_planktivoresA | FishG |
| ***Chromis retrofasciata*** | small planktivores | small_planktivoresA | FishG |
| ***Chromis ternatensis*** | small planktivores | small_planktivoresA | FishI |
| ***Chromis weberi*** | small planktivores | small_planktivoresA | FishQ |
| ***Chromis xanthura*** | small planktivores | small_planktivoresA | FishQ |
| ***Chrysptera rex*** | small planktivores | small_planktivoresA | FishG |
| ***Chrysptera rollandi*** | small planktivores | small_planktivoresC | FishG |
| ***Chrysptera talboti*** | small planktivores | small_planktivoresA | FishQ |
| ***Neopomacentrus azysron*** | small planktivores | small_planktivoresA | FishG |
| ***Pomacentrus brachialis*** | small planktivores | small_planktivoresA | FishI |
| ***Pomacentrus coelestis*** | small planktivores | small_planktivoresA | FishG |
| ***Pomacentrus lepidogenys*** | small planktivores | small_planktivoresC | FishI |
| ***Pomacentrus melanochir*** | small planktivores | small_planktivoresA | FishG |
| ***Pomacentrus moluccensis*** | small planktivores | small_planktivoresA | FishN |
| ***Pomacentrus philippinus*** | small planktivores | small_planktivoresA | FishS |
| ***Acanthurus mata*** | detritivores | detritivores | FishG |

| **Full model** | **Reduced 1** | **Reduced 2** | **Reduced 3** |
|---|---|---|---|
| ***Ctenochaetus striatus*** | detritivores | detritivores | FishS |
| ***Ctenochaetus binotatus*** | detritivores | detritivores | Ctenochaetus binotatus |
| ***Acanthurus lineatus*** | herbivores | herbivoresA | FishS |
| ***Acanthurus nigrofuscus*** | herbivores | herbivoresA | FishJ |
| ***Acanthurus triostegus*** | herbivores | herbivoresB | FishG |
| ***Nasa unicornis*** | herbivores | herbivoresB | FishG |
| ***Zebrasoma scopas*** | herbivores | herbivoresA | FishR |
| ***Cetoscraus bicolor*** | herbivores | herbivoresA | FishV |
| ***Chlorurus gibbus*** | herbivores | herbivoresC | FishK |
| ***Chlorurus sordidus*** | herbivores | herbivoresD | FishH |
| ***Chlorurus bleekeri*** | herbivores | herbivoresD | FishV |
| ***Scarus prasiognathos*** | herbivores | herbivoresC | FishG |
| ***Scarus dubius*** | herbivores | herbivoresE | FishI |
| ***Scarus festivus*** | herbivores | herbivoresE | FishP |
| ***Scarus flavipectoralis*** | herbivores | herbivoresE | FishV |
| ***Scarus forsteni*** | herbivores | herbivoresE | FishQ |
| ***Scarus frenatus*** | herbivores | herbivoresB | FishE |
| ***Scarus globiceps*** | herbivores | herbivoresA | FishM |
| ***Scarus niger*** | herbivores | herbivoresC | FishF |
| ***Scarus psittacus*** | herbivores | herbivoresE | FishV |
| ***Scarus spinus*** | herbivores | herbivoresE | FishH |
| ***Siganus corallinus*** | herbivores | herbivoresD | FishL |
| ***Siganus doliatus*** | herbivores | herbivoresE | FishG |
| ***Siganus punctatus*** | herbivores | herbivoresE | FishO |
| ***Siganus vulpinus*** | herbivores | herbivoresE | FishS |
| **small hebivores** | small hebivores | small_hebivores | FishD |
| **cephalopods** | cephalopods | cephalopods | cephalopods |

| **Full model** | **Reduced 1** | **Reduced 2** | **Reduced 3** |
|---|---|---|---|
| **large invertebrates** | large invertebrates | large_invertebrates | large_invertebrates |
| **small invertebrates** | small invertebrates | small_invertebrates | small_invertebrates |
| **zooplankton** | zooplankton | zooplankton | zooplankton |
| **sponges** | sponges | sponges | sponges |
| **coral/zooxanthallae** | coral/zooxanthallae | coral/zooxanthallae | coral/zooxanthallae |
| **phytoplankton** | phytoplankton | phytoplankton | phytoplankton |
| **epiphilic algae** | epiphilic algae | epiphilic_algae | epiphilic_algae |
| **Detritus** | Detritus | Detritus | Detritus |

*S4: Supplementary results covering all of the threat scenarios and intensities for all functional groups*

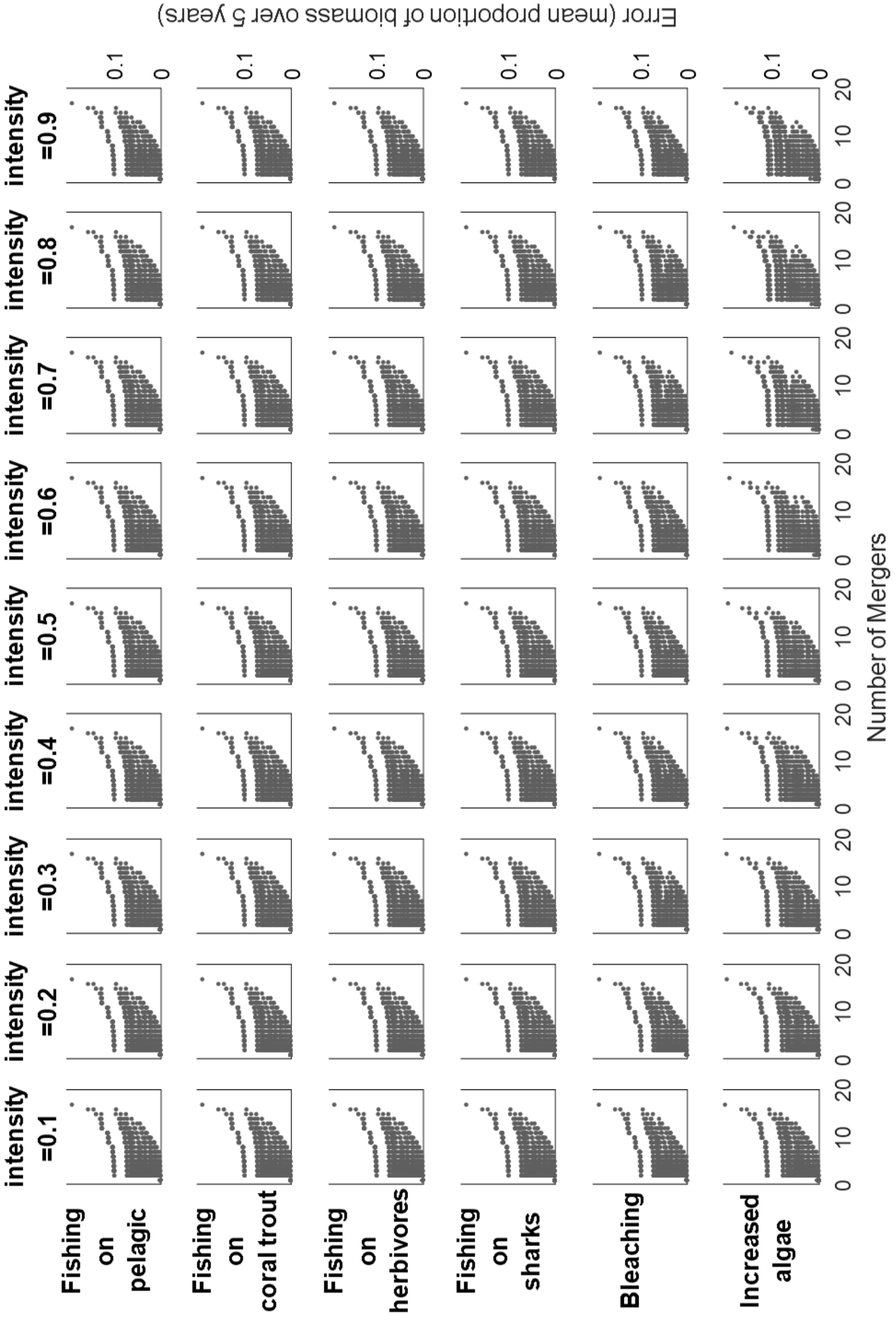


**Figure S4-1:** Error estimates for lutjanids showing all scenarios and intensities

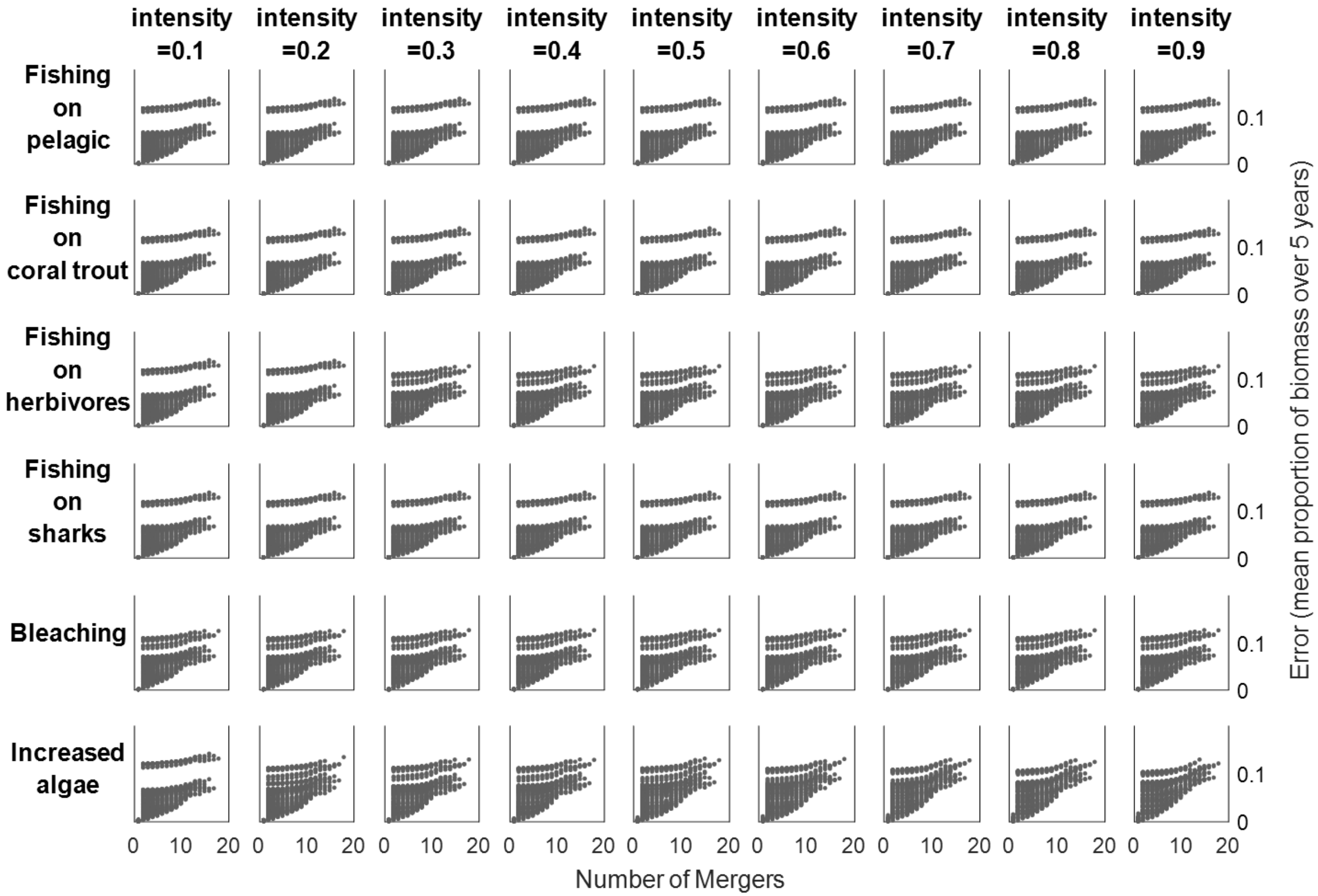


**Figure S4-2:** Error estimates for serranids showing all scenarios and intensities

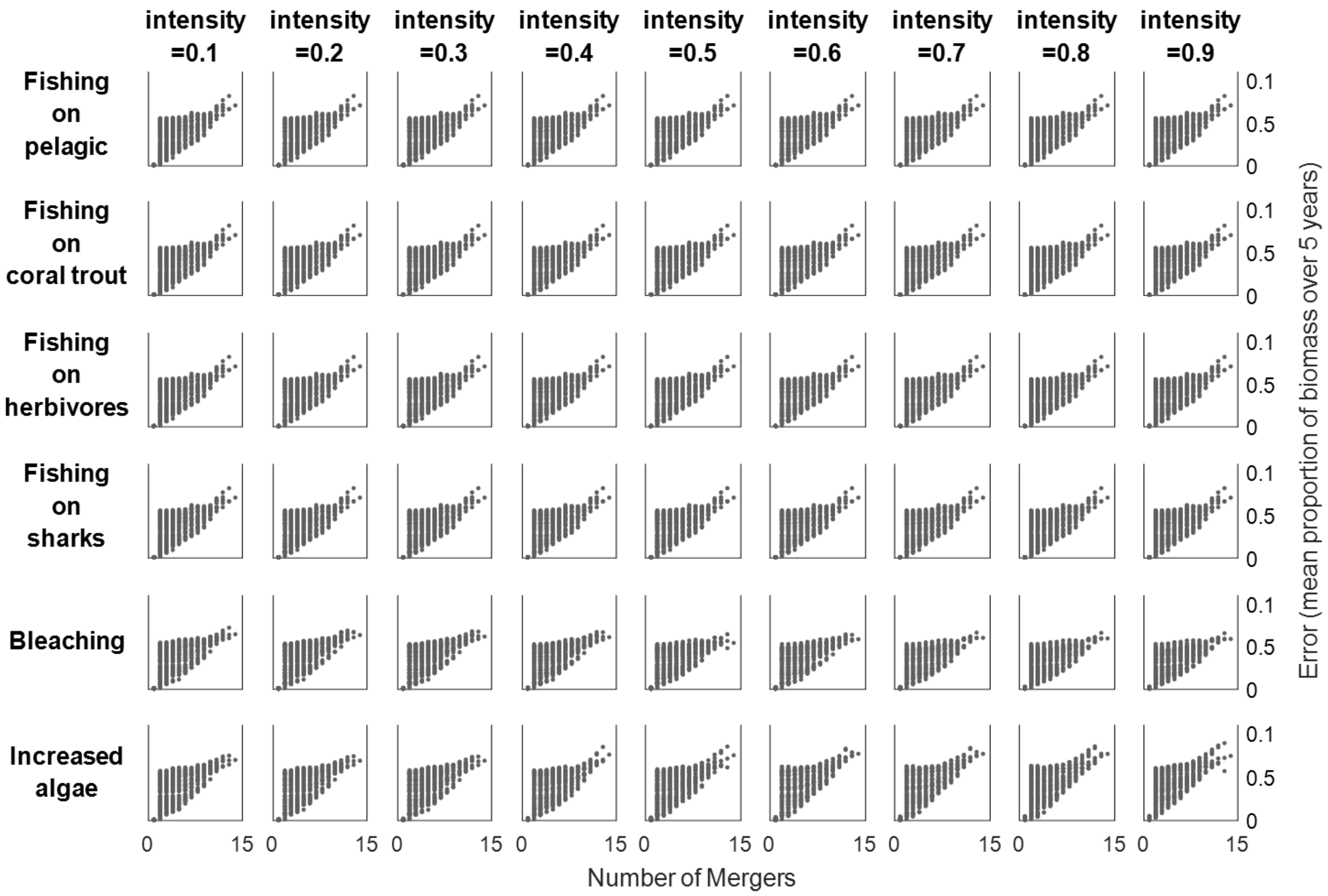


**Figure S4-3:** Error estimates for corallivores showing all scenarios and intensities

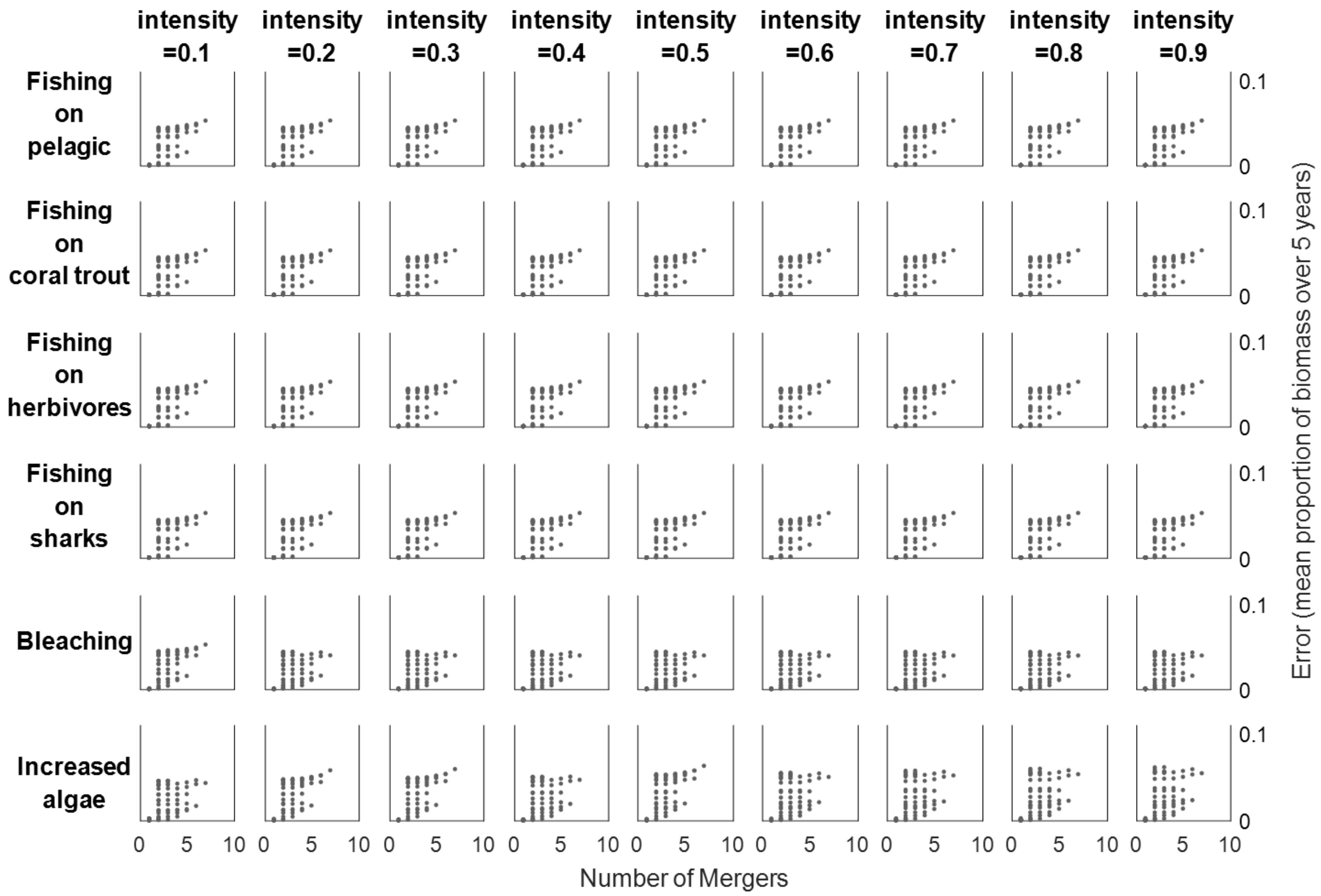


**Figure S4-4:** Error estimates for lethrinids showing all scenarios and intensities

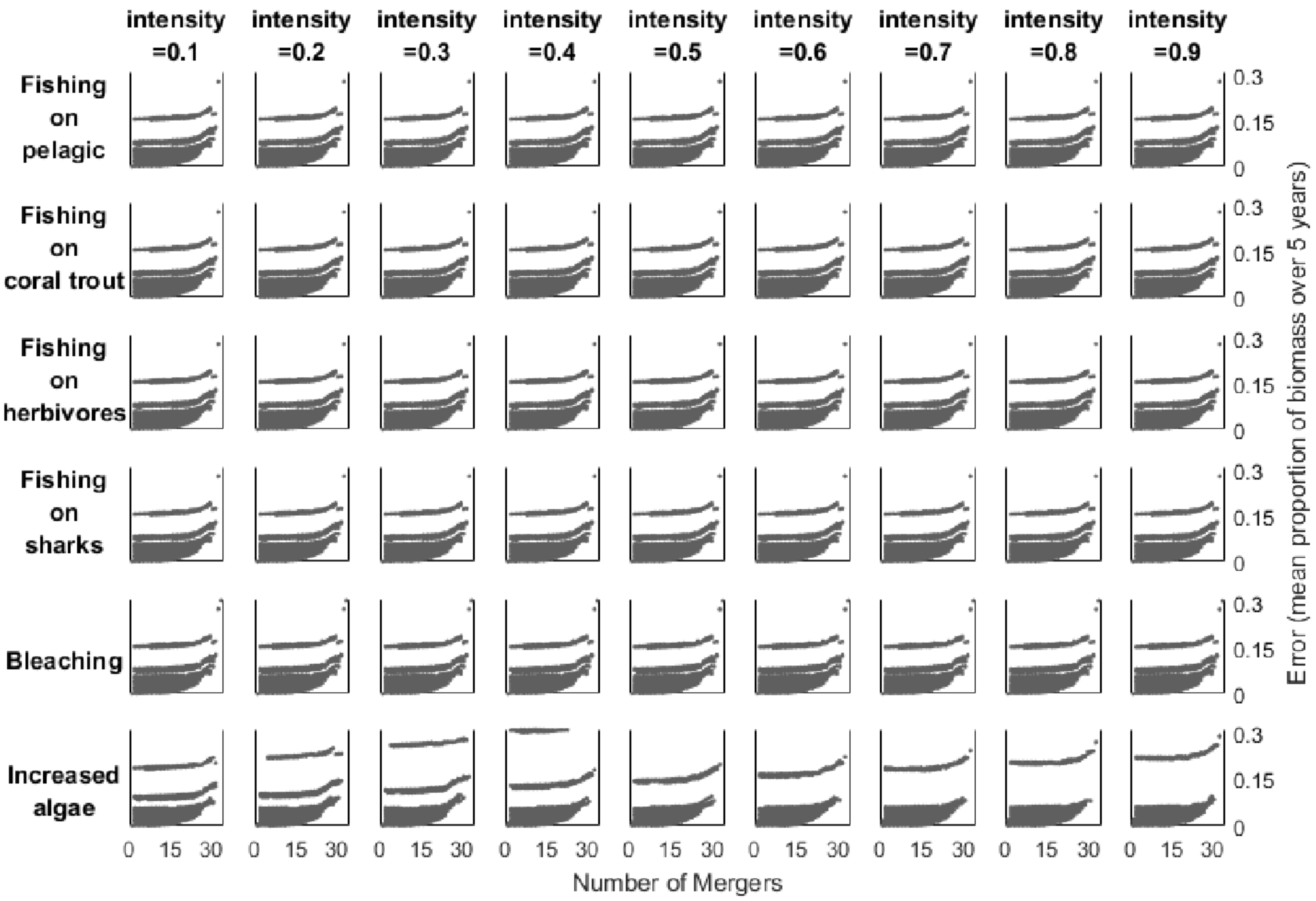


**Figure S4-5:** Error estimates for small planktivores showing all scenarios and intensities

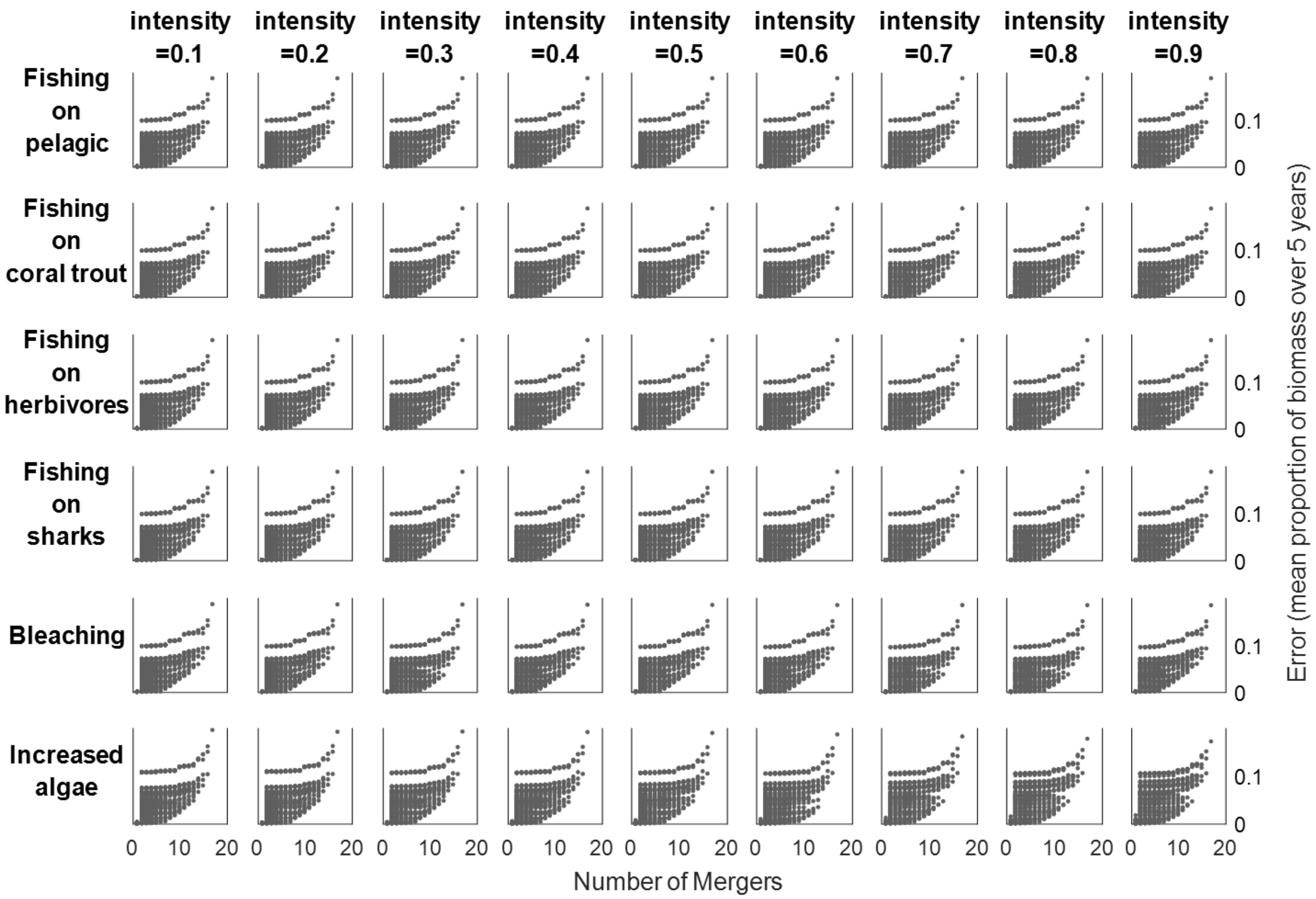


**Figure S4-6:** Error estimates for large planktivores showing all scenarios and intensities

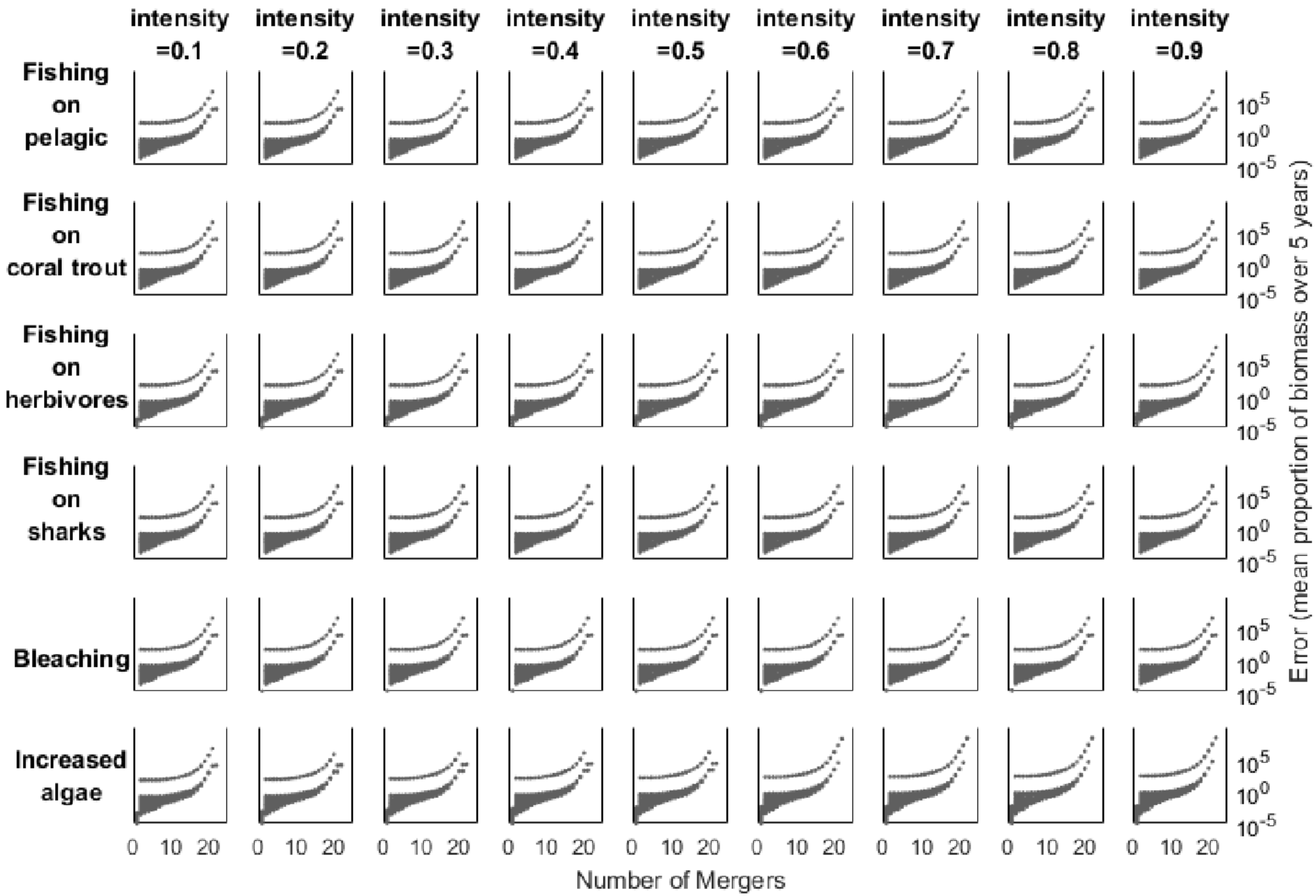


**Figure S4-7:** Error estimates for herbivores showing all scenarios and intensities

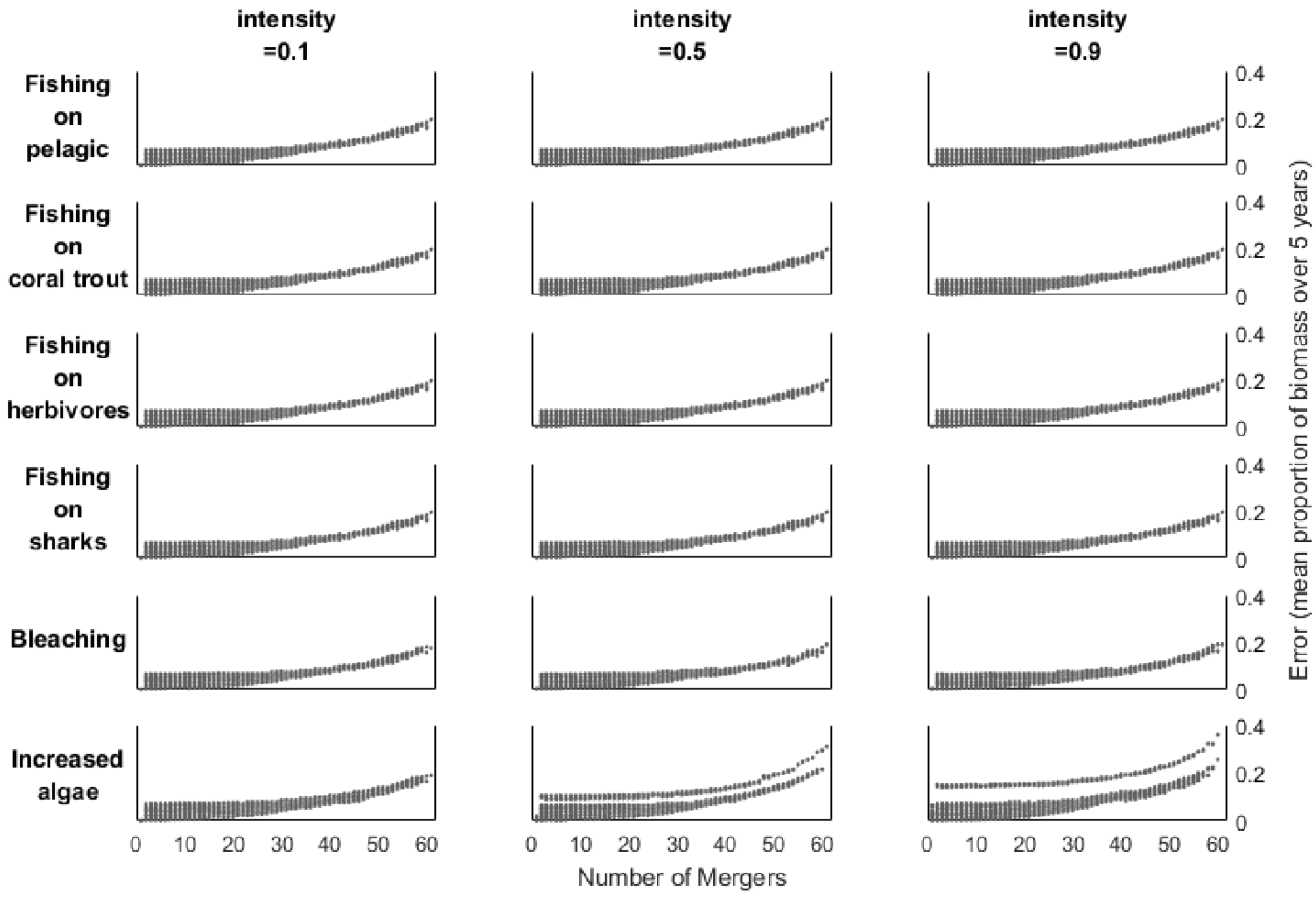


**Figure S4-8:** Error estimates for other demersals showing all scenarios but only three intensities because of computational limitations
